# High-motility pro-tumorigenic monocytes drive macrophage enrichment in the tumor microenvironment

**DOI:** 10.1101/2024.07.16.603739

**Authors:** Wenxuan Du, Bryan Zhou, André Forjaz, Sarah M. Shin, Fan Wu, Ashleigh J. Crawford, Praful R. Nair, Adrian C. Johnston, Hoku West-Foyle, Ashley Tang, Dongjoo Kim, Rong Fan, Ashley L. Kiemen, Pei-Hsun Wu, Jude M. Phillip, Won Jin Ho, David E. Sanin, Denis Wirtz

## Abstract

Enrichment of tumor-associated macrophages (TAMΦs) in the tumor microenvironment correlates with worse clinical outcomes in triple-negative breast cancer (TNBC) patients, prompting the development of therapies to inhibit TAMΦ infiltration. However, the lackluster efficacy of CCL2-based chemotaxis blockade in clinical trials suggests that a new understanding of monocyte/macrophage infiltration may be necessary. Here we demonstrate that random migration, and not only chemotaxis, drives macrophage tumor infiltration. We identified tumor-associated monocytes (TAMos) that display a dramatically enhanced migration capability, induced rapidly by the tumor microenvironment, that drives effective tumor infiltration, in contrast to low-motility differentiated macrophages. TAMo, not TAMΦ, promotes cancer cell proliferation through activation of the MAPK pathway. IL-6 secreted both by cancer cells and TAMo themselves enhances TAMo migration by increasing dendritic protrusion dynamics and myosin-based contractility via the JAK2/STAT3 signaling pathway. Independent from CCL2 mediated chemotaxis, IL-6 driven enhanced migration and pro-proliferative effect of TAMo were validated in a syngeneic TNBC mouse model. Depletion of IL-6 in cancer cells significantly attenuated monocyte infiltration and reversed TAMo-induced cancer cell proliferation. This work reveals the critical role random migration plays in monocyte driven TAMΦ enrichment in a tumor and pinpoints IL-6 as a potential therapeutic target in combination with CCL2 to ameliorate current strategies against TAMΦ infiltration.

## Introduction

Breast cancer is the most frequently diagnosed cancer in women and is the second leading cause of cancer-related deaths^1^. Triple-negative breast cancer (TNBC), defined based on the absence of ER (estrogen), PR (progesterone) and HER-2 expression ^2,3^, is a highly aggressive, invasive, recurrent, and chemotherapy-resistant subtype of breast cancer that additionally features an immunosuppressive tumor microenvironment (TME)^4–6^. Tumor-associated macrophages (TAMΦs) play a pivotal role in orchestrating the immune response in the TME and are found highly enriched in the tumor center and invasive margin, while exerting a broad spectrum of pro-tumoral effects, such as facilitating epithelial-to-mesenchymal (EMT) transition of cancer cells, tissue remodeling, promoting angiogenesis, and mediating immune suppression^7–11^. Since elevated numbers of TAMΦs in TNBC patients is correlated with worse clinical outcomes^12–14^, the inhibition of macrophage recruitment has become one of the main therapeutic targets to address the clinical challenges posed by TNBC^15–17^.

TAMΦs are phenotypically and functionally distinct from tissue-resident macrophages, rapidly expanding via monocyte recruitment during tumor progression^18,19^. Monocytes exhibit high plasticity that enables their differentiation to macrophages and subsequent polarization towards a pro-inflammatory tumor-controlling or a regulatory tumor-promoting phenotype upon exposure to stimuli in TME^20^. Chemokines have long been implicated in macrophage accumulation within tumors^21,22^ and among them, CCL2 induces circulating monocytes to extravasate and, thereafter, modulate differentiated TAMΦ’s chemotaxis in tumor stroma^23^. Anti-CCL2 antibody^24–28^ and CCR2 antagonist^29^ treatments attenuate TAMΦ infiltration in various mouse models with reduced tumor growth and metastasis. However, clinical trials targeting at CCL2/CCR2 axis have proven to be unsuccessful, with limited objective tumor responses observed in patients^16,30,31^.

The current failure of blocking TAMΦ infiltration via inhibiting chemotaxis indicates that our mechanistic understanding of monocyte/macrophage migration in the TME is incomplete. One potentially overlooked consideration in this regard, is the fact that the TME is composed of a highly dense and stiff extracellular matrix (ECM)^32,33^. In solid tumors – including TNB tumors – this tumor-associated collagen-rich stromal matrix functions as a physical barrier and thus requires high migratory potential of immune cells for infiltration^34,35^. Thus, exploring the basal migration of monocytes and macrophages in ECM matrix could allow us to untangle their roles in driving TAMΦ enrichment. Cell migration can be categorized into two distinct patterns: chemotaxis and random basal migration^36^. While chemotaxis drives the directional locomotion of a cell along a chemokine gradient^37^, random migration refers to unbiased cell movement induced by a constant (homogeneous) concentration of soluble factors. Widely adopted *in vitro* cell migration assays for immune cells focus on chemotaxis and do not provide a physiological collagen-rich 3D microenvironment that fully embeds cells^36,38^. Moreover, although the transition of monocytes to macrophages is unidirectional, markers to examine monocytes that recently extravasate into a tissue matrix often overlap with mature macrophage markers, confounding the analysis of these distinct cell types. Consequently, the tissue migration of monocytes before complete differentiation into macrophages has not been extensively studied. Below, we show that the migration patterns of monocytes and macrophages are qualitatively and quantitatively different, a difference that is accentuated in the presence of cancer cells.

In this study, we unraveled the near-complete loss of high basal migration capability of monocytes upon differentiation into macrophages, which implies that macrophages are highly unlikely to actively infiltrate the TME, and their enrichment in the TNBC tumor is caused by the infiltration of monocytes instead. Hence, targeting monocyte migration is key to inhibit TAMΦ enrichment in tumors. High-motility monocytes, which we refer to as tumor-associated monocytes (TAMos), are induced by secreted IL-6 predominantly from cancer cells and regulated via the JAK2/STAT3 signaling pathway in the TME. TAMos are pro-tumoral, as they induce hyper proliferation in TNBC cancer cells by upregulating *FOS*/*JUN* expression to activate the MAPK pathway. After enhanced migration and chemotaxis were demonstrated to be independently modulated by IL-6 and CCL2, we further validated the indispensable role of random migration in preventing monocyte/macrophage infiltration and tumor growth in a syngeneic TNBC 4T1 mouse model. Our work suggests that the combined inhibition of random migration and chemotaxis can greatly ameliorate current therapeutic strategies against TAMΦ recruitment.

## Results

### Monocytes are highly motile compared with differentiated macrophages, a differential motility that is further modulated by the tumor microenvironment

We first characterized the motility of monocytes and macrophages in 3D *in vitro* settings (in the presence/absence of TNBC cells) to reduce potential confounding effects of other cells in the TME. Utilizing a 3D collagen matrix model coupled with time-resolved phase contrast microscopy^39,40^, we tracked the random cell movements of individual macrophages/monocytes and computed migration-related parameters from individual cell trajectories via a customized code (Fig. 1A). To better capture human macrophages/monocytes heterogeneity, which is absent in monocytic cell lines (e.g. THP-1, U937), primary classical human monocytes were isolated from donor peripheral blood mononuclear cells (PBMCs) and differentiated into various phenotypes of macrophages either via a combination of cytokines or tumor conditioned medium, including pro-inflammatory (or M1, with IFN-γ and LPS), regulatory (or M2, with IL-4 and IL-13) and tumor-associated macrophages (Fig. 1A and S1A).

**Fig. 1.**
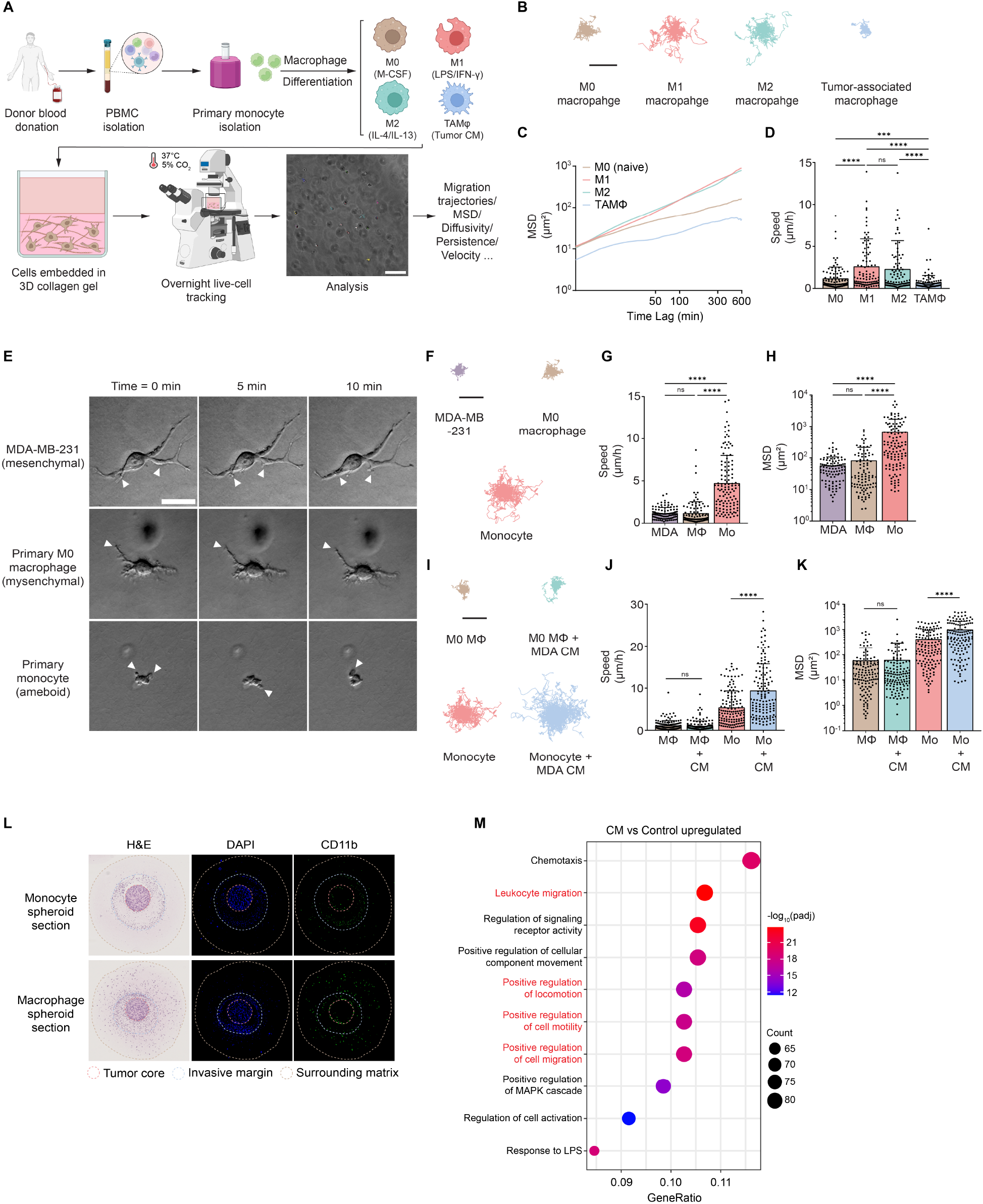
Monocytes exhibit much higher basal random migration capability than macrophages, a difference further enhanced by factors secreted by cancer cells. (**A**) Illustration cartoon of our *in vitro* 3D random migration assay. Primary human monocytes isolated from donor PBMCs via negative selection and differentiated M0 (with M-CSF) / M1 (with LPS/IFN- γ)/ M2 (with IL-4/IL-13) and tumor-associated macrophages (with tumor conditioned medium) were embedded into a 1 mg/mL 3D collagen gel and maintained at 37°C, 5% CO2. Overnight tracking videos were recorded, and individual cells were tracked and analyzed with a customized code for trajectories, mean squared displacement (MSD) and other migration parameters. (**B**) Trajectories of M0/M1/M2 and tumor-associated macrophages during 10 h tracking with a time interval of 5 min. The trajectories of all tracked cells were plotted with the same origin. N= 2 donors, n=50 for a total of 100 cells tracked for each phenotype of macrophages. Scale bar, 150μm. (**C**) Mean MSD of all tracked cells plotted against time lag (min) for each phenotype of macrophages. A smaller slope represents smaller displacements within the same migration time, suggesting lower random migration capability. (**D**) Time-independent speeds during the 10 h tracking were calculated via a customized code and plotted. Each dot represents the speed of an individual cell. (**E**) Representative DIC images obtained with a 20x objective for MDA-MB-231 triple negative breast cancer cell (mesenchymal), naïve M0 macrophage (mesenchymal) and monocyte (ameboid) at 0 min, 5 min and 10 min. Morphological features during active cell migration, including dendritic protrusions and blebs, were marked. Scale bar, 20 μm. (**F-H**) Trajectories, speeds and MSDs of MDA-MB-231 cells, M0 macrophages and monocytes during 6 h tracking (3-min time interval). Monocytes and macrophages shared the same donor. More than 100 individual cells were tacked for each cell type. Scale bar, 250μm. (**I-K**) Trajectories, speeds and MSDs of monocytes and naïve M0 macrophages upon exposure to MDA-MB-231 cancer cell conditioned medium. N= 2 donors, n=50 for a total of 100 cells tracked. Scale bar, 250 μm. (**L**) FFPE sections of 72 h spheroids stained for H&E, DAPI (blue channel) and anti-human CD11b (green channel). Representative scanned images were shown for 1 spheroid for illustration purposes. The section of spheroid was divided into tumor core region, invasive margin region and surrounding matrix region based as annotated. (**M**) Unbiased GO enrichment assay on upregulated genes in MDA CM treated monocytes. Top 10 gene sets with highest gene ratios were plotted and cell migration-related biological processes were labeled red. RNA was extracted from monocytes digested from 3D collagen matrix after 24 h treatment with MDA CM. N=3 biological replicates. For migration assays, characterized speeds, MSDs and motile fractions are plotted as mean + standard deviation. Non-parametric Mann-Whitney test was used for statistical analysis comparing 2 groups (****P≤0.0001, ***P≤0.001, ns P˃0.05). Kruskal-Wallis ANOVA with Dunn’s test was used for statistical analysis for multiple comparisons (****P≤0.0001, ***P≤ 0.001).

As in previous reports^41,42^, upon differentiation from M0 naïve macrophages, M1 macrophages exhibited an elongated morphology with upregulation of co-stimulatory factor CD80, while M2 and tumor-associated macrophages polarized towards a rounded morphology, with upregulation of scavenger receptor CD163 (Fig. S1, A and B). We found that despite differences in morphology and surface marker expression, M1 and M2 macrophages had nearly identical migration capabilities. Both M1 and M2 macrophages showed increased motility compared to their precursor M0 macrophages, evidenced by longer trajectories in 3D collagen matrices (Fig. 1B) and elevated mean squared displacements (MSD - Fig. 1C). M1 macrophages migrated at the average speed of 2.61 μm/h, which is 2.2-fold higher than naïve M0 macrophages that migrated at a speed of 1.17 μm/h (Fig. 1D). Tumor-associated macrophages, on the other hand, showed the lowest motility among all phenotypes (0.69 μm/h, Fig. 1D).

After extracting additional migratory parameters (diffusivity, persistence, anisotropy) from the analysis of the cell trajectories^39^, clustering analysis of the migratory spatial-temporal profile of these different macrophages^43^ revealed a shift in migration mode, from a high diffusivity, high persistence pattern towards a low diffusivity, low persistence pattern in tumor-associated macrophages (Fig. S1C-E). The loss in migration capability of tumor-associated macrophages indicates that the extensive (active) infiltration and recruitment into the tumor microenvironment is not due to mature macrophage migration. Rather, these data suggest that a “migratory phenotype” of macrophages must exist prior to full differentiation and subsequent enrichment in the tumor, which led us to focus on early-stage recruited monocytes.

Reflected light microscopy revealed that naïve M0 macrophage adopted a mesenchymal migration mode similar to breast cancer cell, featuring actin-rich lamellipodia and high protrusion dynamics (Fig. 1E). However, both failed to generate net locomotion within the tracking time of 10 min in the 3D collagen matrix. In contrast, monocytes showed rapid motility driven by pseudopods and hydrostatically generated blebs that resembled ameboid migration^44,45^ (Fig. 1E). Extending the tracking period to 6 h, monocytes covered significantly longer distances (Fig. 1F) and featured a 4-fold increase in speed and 8-fold higher mean squared displacement (MSD) compared with naïve macrophages (Fig. 1, G and H). Exposing monocytes to conditioned medium from MDA-MB-231 TNBC cells (MDA CM) in the collagen matrix (Fig. S1F) further increased cell migration to a speed of 9.41 μm/h (Fig. 1J), an astonishing 8-fold increase compared with macrophages. This difference in migration between monocytes and macrophages was evident across multiple donors (Fig. S1G-I). Further loss in migration capability was induced when naïve M0 macrophages were treated with MDA CM (Fig. 1I-K), similar to what we observed with TAMΦs (Fig. 1D). Notably, the significant migration modulation in monocytes by tumor conditioned medium was rapid (<6 h) and preceded their differentiation into macrophages^46^ (typically 7 days). Indeed, incubating monocytes with MDA CM for up to 7 days completely polarized them into tumor-associated macrophages; they featured a larger cell body and attenuated cell migration capability (Supplementary Video 1).

To further investigate differences in migration between monocytes and differentiated macrophages, a multi-compartment co-culture spheroid system^47,48^ was used where MDA-MB- 231 TNBC cells were embedded at high density in a Matrigel matrix core to mimic a solid tumor with a basement membrane^48^, while monocytes/macrophages were embedded in the outer collagen matrix which mimicked the stromal layer (Fig. S1J). After 72h of co-culture, the enrichment in CD11b^+^ monocytes in the MDA invasive margin and tumor core area was observed, implying that monocytes were recruited and migrated towards the tumor core. In contrast, macrophages, due to their inability to migrate rapidly, stayed evenly distributed in the surrounding matrix, with negligible sign of recruitment into the core (Fig. 1L). In terms of basement membrane invasion, macrophages that initially localized close to the MDA tumor core were also able to infiltrate at a later time point of day 5 (Fig. S1K). These combined results suggest that monocytes, and not fully differentiated macrophages, have a highly motile phenotype, which allows them to move through the dense collagen-rich stromal matrix, which we refer to as “infiltration”. The consequent “enrichment” of tumor-associated macrophages in the tumor microenvironment is therefore a result of this active monocyte infiltration, which is then followed by their differentiation.

The high-motility phenotype of monocytes was not donor-dependent (Fig. S1, G and H) and similar observations were made with monocytic cell lines THP-1 and U937 (Supplementary Fig. S1L-O). This increased motility was caused by a shift in the distribution of monocytes from a low-motility phenotype towards a high-motility phenotype. We define high-motility monocytes as monocytes that show an MSD larger than the square of the averaged monocyte diameter (∼10 μm). We found a higher fraction of monocytes with a high motility phenotype under CM conditions (Fig. S1I). RNA-seq analysis of CM-treated monocyte was well aligned with the above migration results. We found a notable enrichment of biological processes with high gene ratios (4 out of 6 in the 10 gene sets ranked with highest gene ratios) related to cell migration (Fig. 1M). We propose that a high-motility phenotype is characteristic of monocytes conditioned by cancer cells and refer to these cells as “tumor-associated monocytes” (TAMos).

### Tumor-associated monocytes adopt a high-motility phenotype with elevated dendritic protrusion dynamics and contractility via secreted IL-6

We first examined the possibility that the increased motility of TAMos was associated with general cell activation by treating monocytes with a titration of lipopolysaccharides (LPS) following the same timescale as the conditioned medium treatment. Our results showed that LPS exposure exerted little influence on monocyte migration (Supplementary Fig. S2A). Considering the relatively short lifetime of monocytes without colony-stimulating factor (M-CSF), we exposed monocytes to M-CSF and found that cell viability did not contribute to the increased motility of monocytes either (Supplementary Fig. S2B). A previous secretomic assessment showed that MDA-MB-231 cells embedded in 3D collagen matrix selectively secrete high concentrations of cytokines IL-6 and IL-8^49^. When evaluating the clinical outcomes of patients with triple negative breast cancer, the expression of high IL-6 and IL-8 is associated with worse overall survival (Supplementary Fig. S2C). Therefore, we evaluated IL-6 and IL-8 as candidate molecules driving the high-motility phenotype of TAMos.

To assess if IL-6 and IL-8 secreted by MDA-MB-231 cells modulated monocyte motility, we exogenously added each cytokine or their combination to monocytes in 3D collagen matrix (Fig. 2A). While IL-8 alone failed to induce a higher motility, exposure to IL-6 increased monocyte migration speed and motile fraction to levels similar to those observed with MDA CM treatment (Fig. 2B). Flow cytometry indicated a high expression level of IL-6R, but not IL-8R, on CCR2^+^ classical monocytes (Fig. 2C). Interestingly, non-classical monocytes, which predominantly remain in peripheral blood and are not typically recruited to tissues^50^, expressed significantly lower levels of IL-6R compared to classical monocytes (Fig. 2D), which supports our hypothesis that IL- 6 and IL-6R is essential for monocyte tissue migration. We next analyzed a public single-cell atlas of human breast cancers^51^. Among all annotated cell subsets, *IL6R* and *CCR2* were specifically expressed in monocyte and macrophage populations (Fig. 2E).

**Fig. 2.**
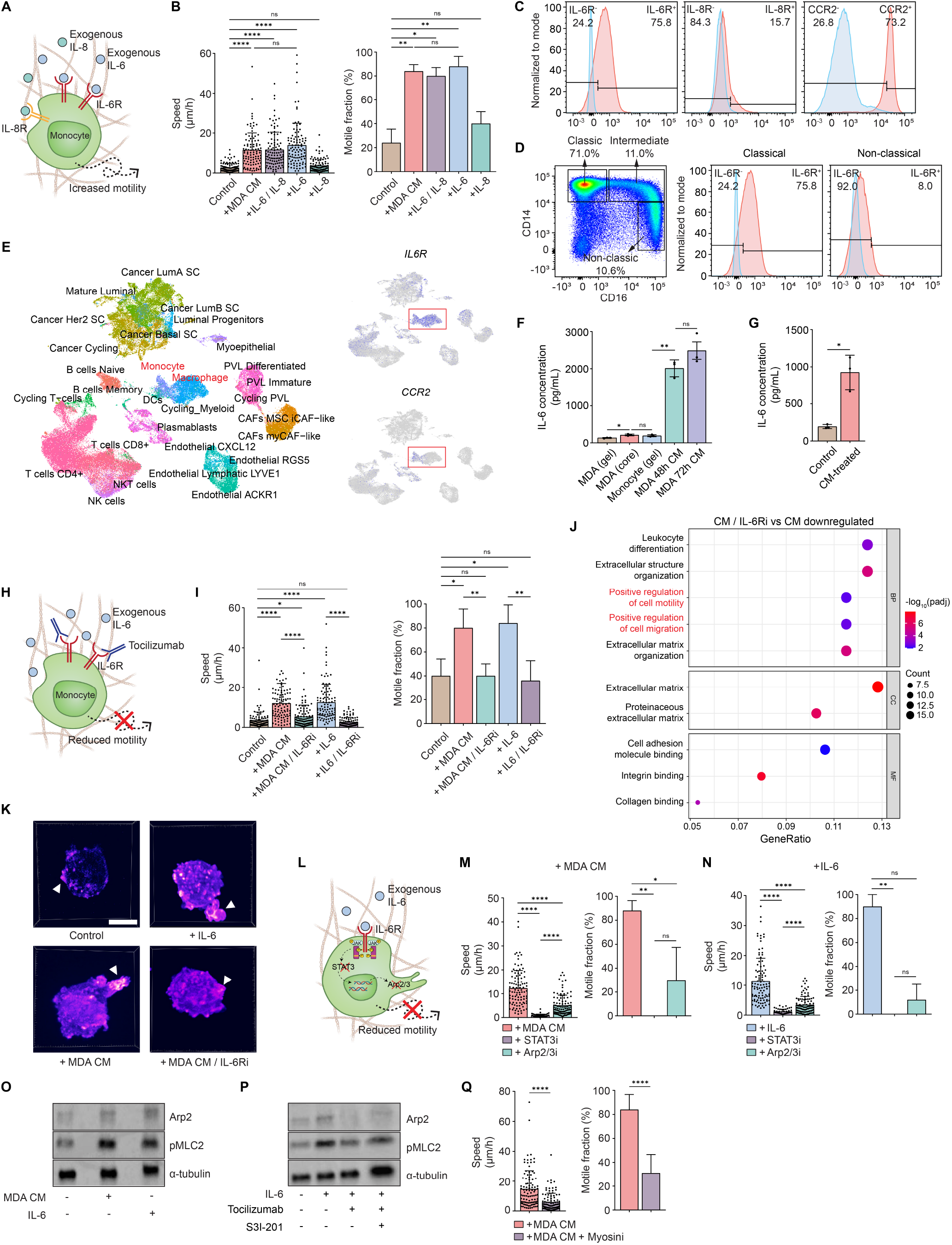
High-motility functional phenotype of TAMo is regulated via an IL-6/JAK2/STAT3 signaling pathway. (**A**) Illustration cartoon of exogeneous cytokine treatment. Monocytes in 3D collagen matrix were incubated for 6 h with 1 ng/mL of IL-6 or IL-8 and then tracked for motility. (**B**) Speeds and motile fractions of monocytes treated with IL-6, IL-8 or IL-6+IL-8. MDA CM- induced TAMo was used as a positive control. (**C**) Surface expressions of IL-6R, IL-8R and CCR2 on classical human monocytes via flow cytometry. Positivity was gated using fluorescence minus one (FMO) control. (**D**) Classical (71.0%), intermediate (11.0%), and non-classical (10.6%) monocytes were gated via CD14/CD16 surface expressions after pan-monocyte isolation. (**E**) Single-cell atlas of human breast cancer with annotated cell subsets (GSE176078) and *IL6R* and *CCR2* expression (high – blue; low – gray). (**F**) Comparison of IL-6 secretion between high-density MDA-MB-231 tumor cores (to mimic tumor microenvironment) and low-density monocytes or MDA-MB-231 cells embedded in 3D collagen gel at 100 cells/mm^3^. N = 3 biological replicates. Unpaired t test with Welch’s correction was used for statistical analysis (**P≤0.01, *P≤0.05, ns P˃0.05). (**G**) IL-6 ELISA from TAMos under indicated conditions. N=3 biological replicates, n = 3 technical replicates. (**H**) Illustration cartoon of functional IL-6R blockade via tocilizumab. 1 ng/mL exogenous IL-6 and 150 nM tocilizumab were added simultaneously to monocytes and incubated for 6 h at 37°C, 5% CO2 before tracking. (**I**) Speeds and motile fractions of monocytes under indicated treatments. (**J**) Top 10 downregulated gene sets in TAMos when IL-6/IL-6R interaction was inhibited. Cell migration related biological processes were labeled red. (**K**) Representative lattice light sheet microscopy images recording active dynamics of cell dendritic protrusions. Scale bar, 5 μm. (**L**) Illustration cartoon showing that IL-6 signaling pathway was inhibited via targeting at downstream transcription factor STAT3 and actin nucleator Arp2/3 complex. (**M-N**) Speeds and motile fractions of monocytes following treatment of 10 μM STAT3 inhibitor S3I-201 or 50 μM Arp2/3 inhibitor CK-666. (**O-P**) Representative western blot results for Arp2 and pMLC2 expression in monocytes treated as indicated. (**Q**) Speeds and motile fractions of TAMos with or without inhibiting myosin II activity using 50 μM blebbistatin. For migration assays, N = 2 donors, n = 50 for a total of 100 individual cells were tracked for each condition in all experiments. Speeds and motile fractions (N = 2 donors, n = 5 technical replicates) are plotted as mean + standard deviation. Non-parametric Mann-Whitney test was used for statistical analysis comparing 2 groups (****P≤0.0001, **P≤0.01, ns P˃0.05). Kruskal-Wallis ANOVA with Dunn’s test was used for statistical analysis comparing groups with control (****P≤0.0001, **P≤0.001, *P≤0.05, ns P˃0.05).

Monocytes can secrete pro-inflammatory cytokines, including IL-6, upon activation in anti- bacterial immunity and inflammation^52,53^. While in the control condition where monocyte-derived IL-6 was not sufficient to promote migration, we still conducted Elisa assay to quantify the secretion of IL-6 from monocytes and MDA-MB-231 cells. MDA-MB-231 cells and monocytes seeded in 3D collagen matrix at same low density showed negligible difference in IL-6 secretion (214 pg/mL for MDA-MB-231 and 194 pg/mL for monocyte, Fig. 2F). However, the conditioned medium that we harvested from solid MDA tumor core, where MDA-MB-231 cells were seeded at higher density to mimic extensive tumor growth, contained significantly higher level of soluble IL- 6, reaching ∼2 ng/mL (Fig. 2F). This indicates that during tumor progression, high levels of IL-6 secreted by cancer cells is essential for the induction of the high-motility phenotype of early-stage recruited monocytes. Interestingly, TAMos increased IL-6 secretion by 4.8-fold to a level of 925 pg/mL (Fig. 2G) and featured upregulated surface IL-6R expression within 6 h of MDA CM treatment (Fig. S2D), which might self-impose these high-motility state.

To show causality and since IL-6 is endogenously produced by both MDA-MB-231 cells and TAMos, we investigated whether functional blockage of IL-6/IL-6R interaction using anti-IL-6R recombinant monoclonal antibody tocilizumab could counteract the increased motility of TAMos induced by IL-6 (Fig. 2H). Blocking IL-6/IL-6R interaction hampered TAMos’ migration and, correspondingly, pushed TAMos into a low-motility state, with a motile fraction dropping to a level similar to control monocytes (Fig. 2I). Unbiased GO enrichment analysis on tocilizumab-treated TAMos showed downregulation of the cell migration-related gene sets that were initially upregulated in TAMos (Fig. 2J and 1M). In sum, IL-6 is necessary and sufficient to induce monocytes towards a high-motility phenotype.

JAK (Janus kinase) and STAT3 (signal transducer and activator of transcription 3) are downstream effectors in the IL-6 signaling pathway^54^. Based on previous work, the JAK2/STAT3 pathway regulates the Arp2/3 complex that nucleates F-actin assembly and mediates protrusions in cancer cells^49^. We tagged monocytic U937 cell with LifeAct-GFP^55^ via lentiviral transduction and monitored its migration in 3D collagen matrix under lattice light sheet microscopy at sub-minute temporal resolution. Generation and active dynamics of dendritic protrusions were visualized in high-motility TAMos induced by MDA CM and IL-6 (Fig. 2K, Supplementary Video 2). The annotated single-cell breast cancer atlas ^51^ also demonstrates high expression levels of *JAK2* (specific to the monocyte/macrophage population) and *STAT3* in subsets of monocyte/macrophage (Fig. S2E). We hypothesized that IL-6 signals activated the JAK/STAT3/Arp2/3 pathway in TAMos to modulate increased motility and subsequently tested the effects of inhibiting STAT3 via S3I-201^56^ and Arp2/3 via CK-666^49^ (Fig. 2L). Exposure to either inhibitor resulted in a marked decrease in TAMo speed and a shift into a low-motility state (Fig. 2, M and N). Clustering analysis of the monocyte migration’s spatial-temporal profile revealed a shift of migration patterns from a high diffusivity, high persistence pattern towards a low diffusivity, low persistence pattern in TAMos upon STAT3/Arp2/3 inhibition (Fig. S2F-H). Finally, Western Blot analysis showed that Arp2 expression was upregulated in TAMos induced by MDA CM and IL-6 (Fig. 2O), while downregulated when the IL-6 pathway was inhibited via either tocilizumab or STAT3 inhibitor S3I-201 (Fig. 2P).

Adopting an ameboid migration mode (Fig. 1E) indicates that monocyte migration in 3D matrix is adhesion-independent and instead modulated by cell contractility that drives cell “squeezing” through the collagen fibrous network^45,57,58^ (Fig. S2I). Monocyte motility was negatively correlated with collagen concentration and associated smaller pore size of the collagen matrix^59^ (Fig. S2J). We evaluated the expression of phosphorylated myosin light chain II in TAMo to examine if this cytoskeletal component was downstream of the IL/6/JAK2/STAT3/Arp2/3 pathway. Like Arp2, pMLC2 (phosphorylated myosin light chain 2) was increased in TAMos and decreased upon inhibition of the IL-6 pathway (Fig. 2, O and P). Functionally blocking myosin II activity via blebbistatin also hampered TAMo migration (Fig. 2Q). In sum, these results indicate that IL-6 enhances monocyte dendritic protrusion dynamics and contractility via a JAK2/STAT3/Arp2/3 signaling pathway to induce high-motility TAMos.

### Pre-differentiated tumor-associated monocytes promote proliferation of cancer cells

Derived from recruited monocytes and re-educated by the tumor microenvironment, tumor-associated macrophages have been well characterized for their pro-tumoral effects, including promoting cancer cell invasion/metastasis via tissue remodeling and orchestrating immunosuppression^11^. However, the transition from monocyte to macrophage requires days of differentiation and little is known on the role that early-stage “pre-differentiated” monocyte may play in tumor progression. Based on our 3D migration results and corresponding RNA-seq data, high-motility TAMos could be rapidly (∼6 h) induced via IL-6 secretion from cancer cells leading to enhanced random migration and subsequent significant monocyte recruitment towards the tumor invasive margin and core within 3 days (Fig. 1L). Consequently, we decided to explore if TAMos could impact tumor growth prior to complete differentiation into low-motility macrophages.

To assess the potential impact of TAMos on cancer cell behavior, we embedded either control polystyrene beads, naïve M0 macrophages or isolated monocytes in the outer collagen layer, respectively, in the multi-compartment spheroid system. With monocytes in the stromal layer, phase-contrast microscopy revealed a noticeably larger expansion of the tumor core, accompanied by more invading breast cancer cells at the invasive margin (Fig. 3A). We applied the PrestoBlue viability assay^47^ to quantitatively monitor cell proliferation in 3D. The relative consistent PrestoBlue signal readings from monocytes and macrophages embedded in the spheroid validated that the increase in signal came mainly from the proliferation of cancer cells in the co-culture system (Fig. 3B). In striking contrast to naïve M0 macrophages which significantly hindered proliferation of cancer cells, monocytes promoted cancer cell proliferation (Fig. 3C). To further explore the ability of monocyte/macrophage to regulate cancer cell proliferation and refine our readouts to account for both proliferation and cell viability, we measured bioluminescence from luciferase-tagged MDA-MB-231 cells (Fig. 3D)^48^. The versatility of the multi-compartment 3D co-culture system also allowed us to adjust the initial spatial distribution of monocytes to be either in the collagen matrix (“matrix co-culture”) or in the core (“core co-culture”) to investigate if migration affects the pro-tumoral effect of TAMos. Consistent with the PrestoBlue assay, TAMos induced proliferation of MDA-MB-231 cells, while naïve M0 macrophages hampered their proliferation. Tumor-associated macrophages hardly exhibited any proliferation-promoting effect (Fig. 3E). This pro-proliferation effect, unique to TAMos, highlights the importance of early-stage recruitment monocytes and can potentially couple with matrix remodeling effect induced by late-stage differentiated tumor-associated macrophages to facilitate tumor dissemination^9,10^. Moreover, monocyte-induced proliferation was most evident when monocytes were seeded together with MDA-MB-231 cells in the tumor core (Fig. 3F), compared to seeding in the collagen matrix. This indicates that TAMos maximize their pro-tumoral effect after extensive migration towards the tumor core, which rationalizes the induction of high motility by the tumor microenvironment.

**Fig. 3.**
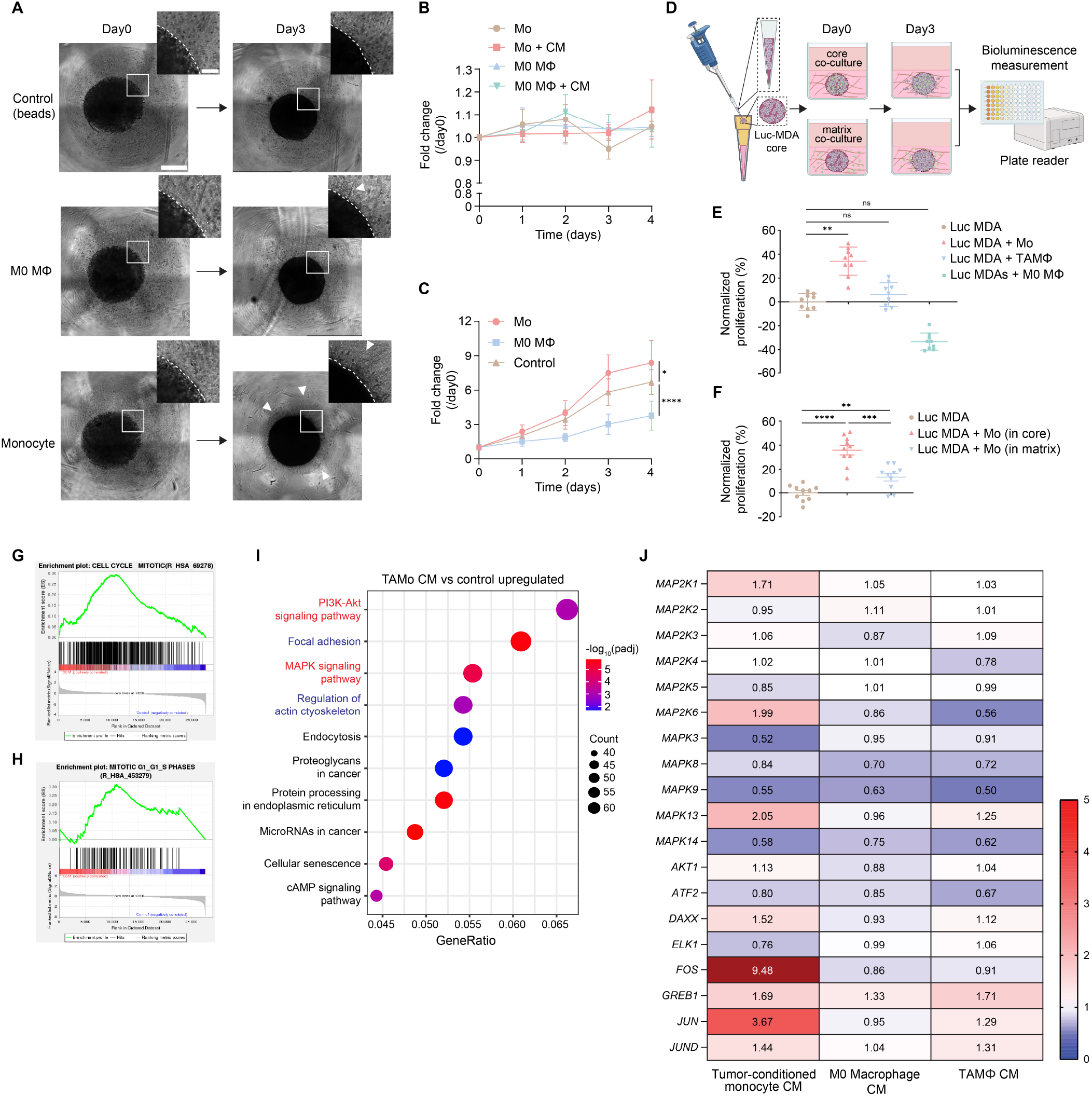
TAMos are pro-tumorigenic before full differentiation into macrophages. (**A**) Representative phase contrast images taken with a 4x objective, and corresponding (**B-C**) PrestoBlue signal intensity, for multi-compartment MDA-MB-231 spheroids co-cultured with beads, M0 naïve macrophages and monocytes. Scale bar, 300 μm (inset, scale bar, 30 μm). N= 3 biological replicates, n=10 technical replicates. Data are plotted as mean ± standard deviation. (**D-F**) Cartoon with experimental set-up and quantification of 3D luciferase assay for MDA-MB- 231 cell proliferation under indicated co-culture systems. Specific proliferation was normalized to control where MDA-MB-231 cells were cultured alone. N = 2-3 donors, n = 3-5 technical replicates. (**G-I**) Pathway enrichment analysis from RNA-seq of MDA-MB-231 spheroids treated with TAMos’ conditioned medium. N = 3 biological replicates. (**J**) Heatmap of fold changes for key genes in MAPK signaling pathway assessed with RT-qPCR from MDA spheroids treated as indicated. N = 3 biological replicates. n = 3 technical replicates. Data for PrestoBlue and Luciferase assay are plotted as mean ± standard deviation. Non-parametric Mann-Whitney test was used for statistical analysis comparing 2 groups (****P≤0.0001, ***P≤0.001, **P≤0.01, *P≤0.05, ns P˃0.05). Kruskal- Wallis ANOVA with Dunn’s test was used for statistical analysis comparing groups with control (**P≤0.001, ns P˃0.05).

We next sought to identify the mechanism by which monocytes could promote tumor growth and hypothesized that these TAMos secrete proteins that induced cancer cell proliferation. Indeed, we observed that monocytes induced MDA-MB-231 proliferation in both the “matrix co-culture” system, where extensive intercellular contact is initially absent, and in the “core co-culture” setting. Consequently, we exposed MDA-MB-231 spheroid with TAMos’ conditioned medium for 48 h and carried out RNA-seq analysis to identify differentially expressed genes in cancer cells. Gene set enrichment analysis of Reactome pathways showed significant upregulation of mitotic cell cycle-related gene sets (Fig. 3, G and H), which reflected the observed high proliferation of breast cancer cells. KEGG pathway analysis revealed the upregulation of PI3K-Akt and MAPK signaling pathways that play critical roles in oncogenesis (Fig. 3I). Previous studies have shown a correlation between activation of ERK and proliferation of cancer cells in multiple models of breast cancer^60^. We performed RT-qPCR analysis on genes associated with the MAPK pathway and our results showed that gene expression of downstream transcription factor *FOS* in the ERK cascade was increased by 9.48-fold and *JUN* in the JNK cascade was concurrently increased by 3.67-fold in TAMo CM-treated MDA-MB-231 cells compared with control cells. These striking results were not observed in breast cancer cells treated with CM from naïve M0 macrophages or tumor-associated macrophages (Fig. 3J).

Collectively, our data demonstrate that TAMos induce cancer cell proliferation by activating the MAPK signaling pathway as a result of factors secreted by these monocytes.

### Random migration and chemotaxis of monocytes are regulated independently by cancer cells via IL-6 and CCL2

The term migration has long been used ambiguously to describe chemotaxis in studies using the traditional transwell assay^36^. Comparably little attention has been devoted to understanding random basal migration of immune cells. Harnessing the 3D collagen gel model, we have investigated how IL-6 in the breast cancer tumor microenvironment enhances monocyte random migration. We next sought to demonstrate how random migration of monocytes played an indispensable role in its recruitment cascade, independently of chemotaxis.

Since CCL2 is the major chemokine that is highly secreted by breast cancer cells and recruits monocyte/macrophage towards breast tumor^23^, we first evaluated if CCL2 impacted monocyte random migration by adding exogenous CCL2 to monocytes in 3D collagen matrix (Fig. 4A). Unlike IL-6, CCL2-treated monocyte exhibited no change in speed and motile fraction compared to untreated cells (Fig. 4B). We then switched to a transwell assay where we seeded monocytes on top of a collagen-coated porous membrane and added CCL2 and IL-6 separately in the bottom chamber (Fig. 4C). In contrast to results from the 3D random migration assay, CCL2, and not IL- 6, induced chemotaxis and recruited more monocytes via transmigration through the porous membrane (Fig. 4D). Conditioned medium from MDA-MB-231 cells, which contains both CCL2 and IL-6, could induce chemotaxis and enhanced random migration simultaneously (Fig. 4D and 1J). Likewise, in a multi-compartment spheroid, we found that monocytes in the collagen stomal matrix were recruited towards the tumor core via both CCL2-mediated chemotaxis and IL-6- mediated enhanced migration (Fig. 4E-G). We tracked the trajectories of monocytes and characterized their directionality either as chemotaxis towards the tumor core (downward) or as non-chemotaxis away from the tumor core (upward). When CCL2-CCR2 interaction was blocked via CCR2 antagonist INCB3344, monocytes lost their chemotactic capability with a biasing ratio < 1 yet kept a high motility phenotype with long trajectories (Fig. 4, F and G). In such a scenario, TAMos with inhibited chemotaxis are still prone to infiltrate tumor given their preserved high motility. When anti-IL-6R antibody tocilizumab was added, monocytes displayed severely reduced migration, as evidenced by the short trajectories, but maintained chemotaxis with a biasing ratio of 2.0. Far from the MDA tumor core, monocyte migration was significantly reduced with neither enhanced migration nor chemotaxis present. We therefore conclude that the recruitment cascade of monocytes combines enhanced random migration together with chemotaxis, one modulated by IL-6, which controls cell speed, while the other modulated independently by CCL2, which controls cell directionality (Fig. 4H). Critically, these results indicate that only by blocking both random migration and chemotaxis can we significantly reduce monocyte recruitment and eventually inhibit tumor-associated macrophage infiltration. Previously we demonstrated that TAMo infiltration induces proliferation of tumor cells (Fig. 3F). Here, combined blockade of chemotaxis and random migration hampered this hyper-proliferation induction in breast cancer cells to a greater extent than by inhibition of either process separately (Fig. 4I).

**Fig. 4.**
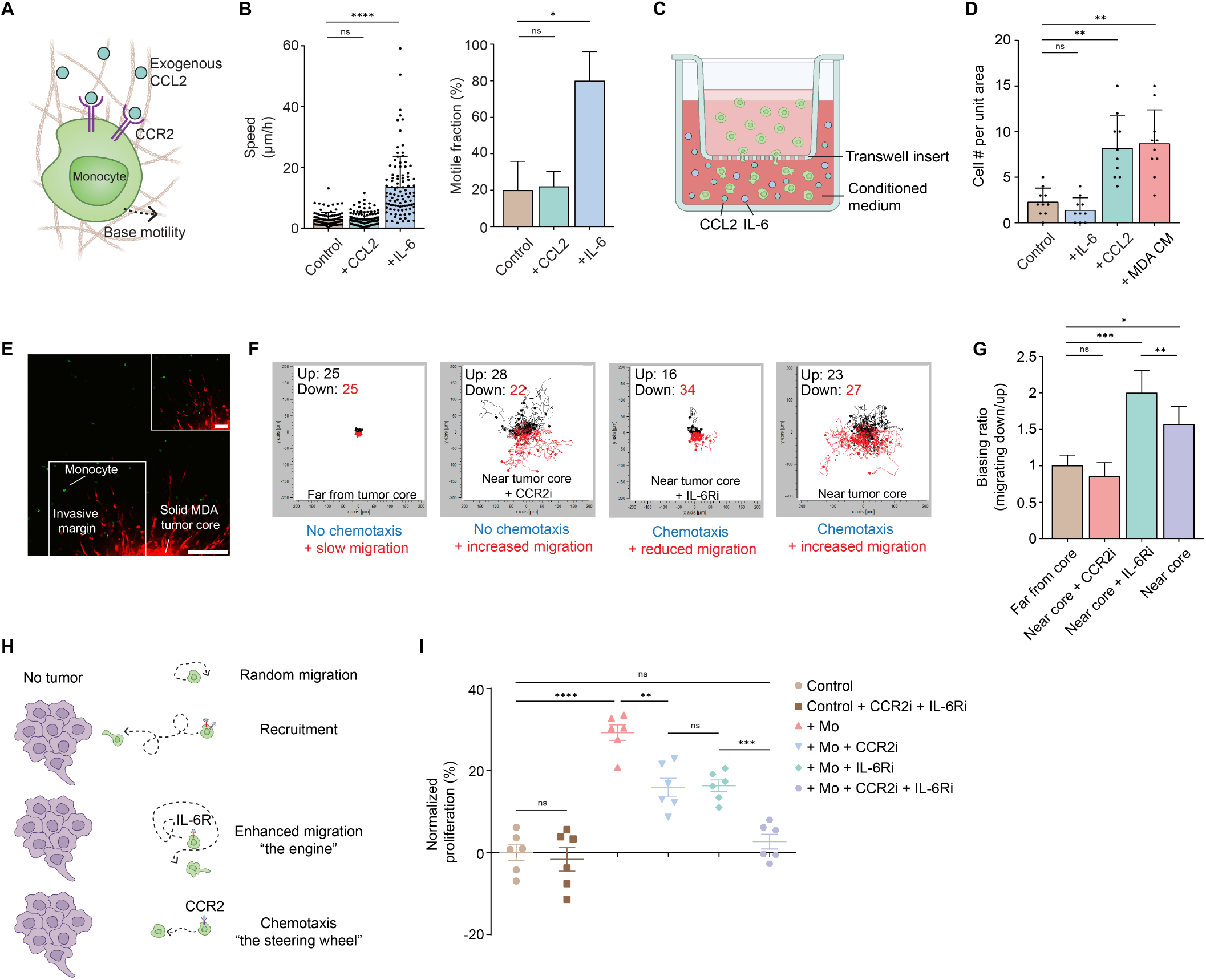
Monocyte recruitment consists of IL-6-mediated enhanced basal migration and CCL2-mediated chemotaxis. (**A**) Primary human monocytes in 3D collagen matrix were treated with chemokine CCL2 exogenously added at 1 ng/mL. (**B**) Characterized speeds and motile fractions of monocytes treated with exogenous CCL2 or IL-6. N=2 donors, n = 50 cells for a total of 100 individual cells were tracked. (**C**) Illustration cartoon of the transwell assay. CCL2 or IL-6 was added to the bottom chamber at 1 ng/mL, respectively. (**D**) Elevated numbers of transmigrated monocytes were counted per unit area (250 μm x 250 μm) in bottom chambers containing MDA CM or CCL2 instead of IL-6. N = 2 donors, n = 5 technical replicates. (**E**) Representative IF image of a multi-compartment spheroid. MDA-MB-231 cells were labeled with CMTPX (red channel) and monocytes were labeled with CFDA-SE (green channel). Scale bar, 300 μm (inset, scale bar, 100 μm). (**F**) Plotted trajectories of monocytes (n = 50 individual cells from 1 technical replicate were shown in each figure) that were close to the invasive margin in each condition. Chemotaxis was inhibited via 200nM of CCR2 antagonist INCB3344 and random migration was inhibited via 150nM of tocilizumab. Monocytes were tracked for 6 h at a time interval of 3 min. Trajectories of monocytes migrating down were labeled red and migrating up were labeled black. (**G**) Biasing ratios reflected the directionality of monocyte migration. A biasing ratio greater than 1 suggested chemotaxis towards tumor core. N = 3 biological replicates, n = 3 technical replicates. Data are plotted as mean + standard deviation. (**H**) Illustration cartoon of monocyte recruitment in the tumor microenvironment. (**I**) Proliferation percentages of MDA-MB-231 breast cancer cells normalized to control after 72 h matrix co-culture of TAMos under different migratory conditions. Data are plotted as mean ± SEM. N=2 donors, n=3 technical replicates. Non-parametric Mann-Whitney test was used for statistical analysis comparing two groups (***P≤0.001, **P≤0.01, ns P˃0.05). Kruskal-Wallis ANOVA with Dunn’s test was used for statistical analysis for multiple comparisons (****P≤0.0001, ***P≤0.001, **P≤0.001, ns P˃0.05).

In addition to driving chemotaxis at tissue level, CCL2 also leads to monocyte extravasation via trans-endothelial migration, which potentially drives the accumulation of monocytes to the stroma- vasculature interface^36,61^. Recent work has shown that cell density can modulate the migration of T cells and cancer cells^35,49^ and we thus explored the possibility that a high density of monocytes could modulate their own migration (Fig. S3A). We found that at higher density, the speed increased up to 4.4-fold and monocytes were 100% motile (Fig. S3B). Even at high density, intercellular collisions were not observed, so we hypothesized that this density-induced motility increase was elicited by monocyte-secreted molecules. To test this hypothesis, we transferred the conditioned medium from high-density (HD) collagen gel (= gel containing a high density of monocytes) onto the low-density (LD) gel and found that it was sufficient to induce a similar increase in migration (Fig. S3B).

A secretomic assay identified IL-6, IL-7, IL-8, IL-17A and CCL2 as secreted by HD monocytes, while undetectable for LD monocytes (Fig. S3C). IL-8 and CCL2 have already been shown to have no effect on monocyte random migration (Fig. 2E and 4B). Among the remaining three factors, only IL-6 recapitulated the enhanced migration phenomenon (Fig. S3D). These findings confirm that IL-6 secreted by breast cancer cells and monocytes themselves at local high density, plays a central role in monocyte migration (Fig. S3E).

### IL-6 depletion in cancer cells prevents monocyte infiltration and reverses cancer cell hyper-proliferation in TNBC syngeneic mouse model

To validate the above high-motility pro-tumorigenic TAMo phenotype and assess the importance of random migration in a more complex *in vivo* onco-immune landscape, we transduced 4T1 mouse triple-negative breast cancer cells with siRNA-expressing lentiviral particle to knock down IL-6 (Fig. S4A) and evaluated the corresponding monocyte/macrophage infiltration in the primary tumor at early (when tumor volume reached half of the sacrificing threshold volume) and late stage (end of study) with blockade of either random migration or chemotaxis in a 4T1 syngeneic mouse model (Fig. 5A). As IL-6 can regulate tumorigenesis^62–64^, we carried out the PrestoBlue assay and showed no difference between the growth of Scr (scramble control) and IL-6 KD (knock down) 4T1 cells in 3D spheroid (Fig. S4B). We then examined the ability of IL-6 sufficient and deficient 4T1 cancer cells to induce the high-motility functional phenotype of mouse monocytes we previously observed with human monocytes (Fig. S4C). Conditioned medium from 4T1 Scr cells increased the random migration of mouse bone marrow monocytes, while conditioned medium from IL-6 KD 4T1 cells did not (Fig. S4, D and E).

**Fig. 5.**
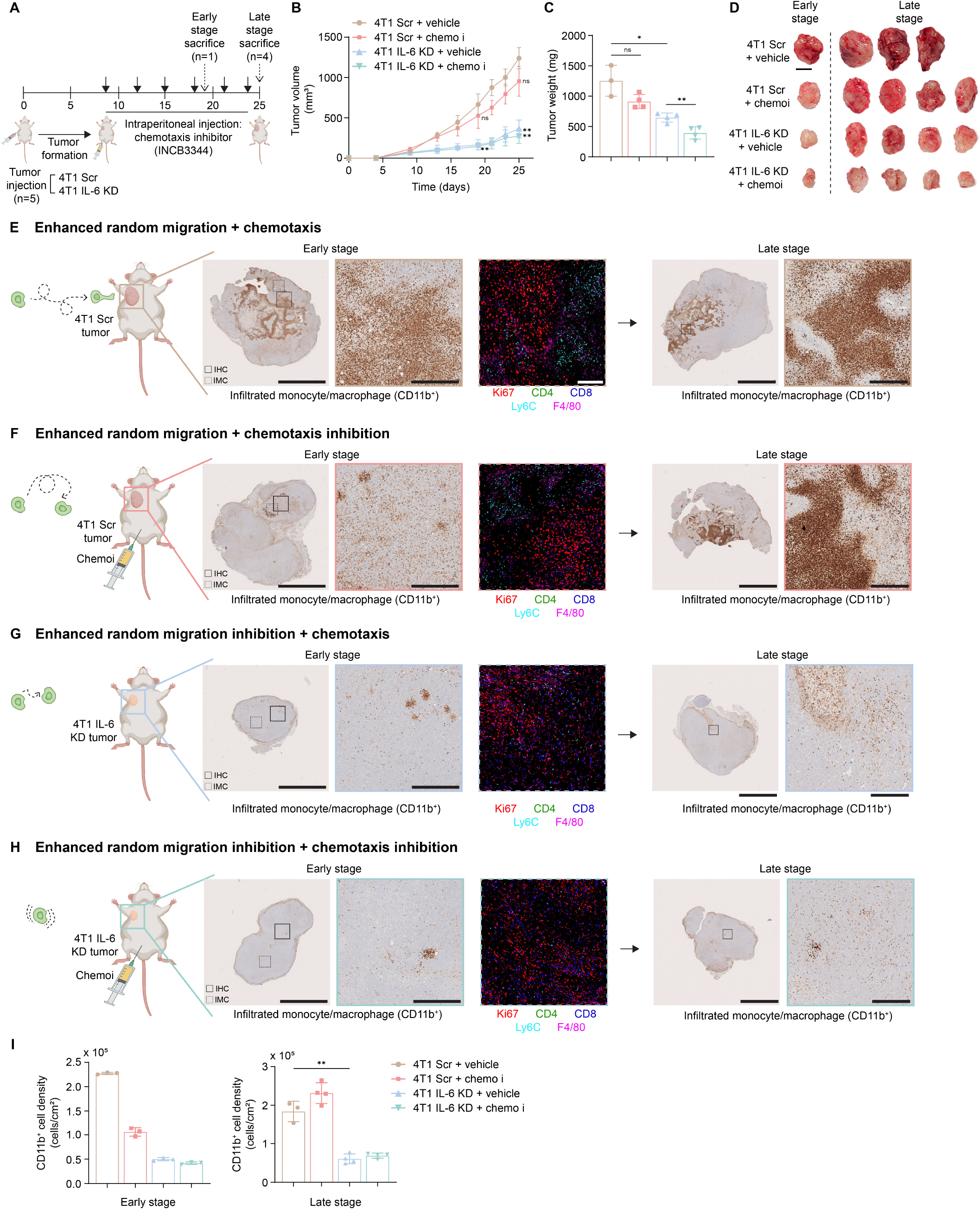
Inhibition of random migration and chemotaxis hampers monocyte infiltration and its pro-proliferation effect *in vivo*. (**A**) Schematic of 4T1 triple negative breast cancer syngeneic mouse model. 4T1 Scr and IL-6 KD cells were orthotopically injected in mammary fat pads of BALB/c mice. Murine CCR2 antagonist INCB3344 was intraperitoneally administered twice a week after palpable tumors were observed. n = 5 mice per group for a total of 4 groups. One mouse per group was sacrificed at early stage (day 19) and four others were sacrificed at end of study (late stage, day 25). (**B**) Time-dependent volumes of primary tumors assessed using calipers for 4 groups. Data are plotted as mean ± standard deviation. (**C**) Tumor weight measurements of harvested tumors for 4 groups at end of study. n = 3 for 4T1 Scr + vehicle group and n = 4 for other 3 groups. Data are plotted as mean ± standard deviation. Unpaired t test with Welch’s correction was used for statistical analysis (**P≤0.01, *P≤0.05, ns P˃0.05). (**D**) Photos for primary tumors harvested at early stage and late stage for four groups. Scale bar, 500 mm. (**E-H**) Tumors from indicated groups were harvested, formalin-fixed-paraffin-embedded and sectioned. Slides were prepared for IHC staining with indicated antibodies. Representative scanned IHC images for early- and late-stage tumors are presented in left and right panels, respectively. Scale bar, 5 mm (ROI scale bar, 500 μm). Representative multicolor IMC images (1000 μm x 1000 μm) are presented in the middle panel. Scale bar, 200 μm. (**I**) Quantitative assessments of CD11b+ monocyte/macrophage density in early- and late-stage tumors. For early stage, three non-consecutive slides from one tumor were IHC stained, counted and plotted per treatment group. For late stage, n = 4 mice per group, except for 4T1 Scr + vehicle group for which n = 3. Unpaired t test with Welch’s correction was used for statistical analysis of domain numbers (**P≤0.01).

In line with our previous observations with human monocytes and MDA-MB-231 cells (Fig.3F and 4I), our in vivo murine tumor models demonstrated that IL-6 production by cancer cells greatly enhances tumor growth, presumably through enhanced monocyte recruitment (Fig. 5B-D). Moreover, as in unsuccessful clinical trials, blocking monocyte chemotaxis alone had a marginal effect on 4T1 tumor growth (Fig. 5B-D). Blocking monocyte random migration by removing IL-6 production significantly reduced tumor volume. Finally, combined inhibition of random migration and chemotaxis resulted in the smallest tumors (Fig. 5B-D), highlighting the potential therapeutic benefit of this combined strategy. To demonstrate that the IL-6 mediated effect on tumor growth was associated with monocyte recruitment, we examined the prevalence of these cells in tumor bearing mice. As monocytes recruited to the tumor microenvironment are differentiated into macrophages (and no single IHC marker can accurately distinguish between monocytes and macrophages), we selected the pan-monocyte/macrophage marker CD11b to assess the infiltration of monocytes in general under different migration conditions. We observed that at early stages of tumor growth, monocytes infiltrated the entire tumor, edge and core, in large amounts in IL-6 sufficient tumor-bearing mice (Fig. 5E, left panel), with continuous prevalence of these several days later (Fig. 5E, right panel). Consistent with the tumor growth trend, inhibition of chemotaxis partly mitigated monocyte infiltration (Fig. 5F, left panel), however this inhibition was transient with later stage tumors displaying comparable numbers of cells (Fig. 5F, right panel). Critically, inhibition of random migration or the combination of the two completely excluded monocytes from the tumor core during both early and late tumor development (Fig. 5, G and H). Interestingly, we detected CD11b staining in these tumors in a “ring” like structure around the tumor edge (Fig. 5, G and H), which might indicate that macrophages already residing in the tissue and preceding tumor growth are excluded from tumors and have poor migratory potential, in line with our previous observations (Fig. 1D). Quantitative assessments of monocyte infiltration were obtained at single-cell resolution using the customized code CODA^65^, validating the trend shown in representative IHC images (Fig. 5I). To assess the infiltration level of all leukocytes, we stained tumors with common leukocyte antigen CD45 and found no difference with or without inhibition of monocyte migration (Fig. S4F), indicating that targeting IL-6 and CCL2 predominantly impacts monocyte infiltration.

To further characterize the immune landscape of the tumor, we performed multiplexing imaging mass cytometry (IMC) analysis^66^. Infiltrated CD11b^+^Ly6C^+^ monocytes were found expressing high level of S100A9 suggested by the co-localization of three markers (Fig. S4G). Fewer Ly6C^+^ monocytes and F4/80^+^ macrophages infiltrated the core of IL-6 deficient tumors, which also displayed increased infiltration of cytotoxic CD8+ T cells, which correlated well with the decreased size of the tumor (Fig. 5E-H, middle panel).

Taken together, we validated the IL-6-mediated enhanced random migration and pro-proliferative effect of TAMos in a 4T1 syngeneic mouse model. Our data show that IL-6 depletion in cancer cells prevents monocyte/macrophage infiltration and reverses cancer cell hyper-proliferation.

## Discussion

Tumor-associated macrophages (TAMΦs) play a critical role in tumor progression^9,10,16^, yet TAMΦs display an extremely limited ability to migrate compared to monocytes and other macrophage phenotypes (Fig. 1D and G). Hence, using triple-negative breast cancer (TNBC), we sought to understand how TAMΦs become enriched in the tumor microenvironment. Our results suggest that monocytes exposed to factors produced by breast cancer cells adopt a highly motile phenotype, which we referred to as tumor-associated monocytes (TAMos). In contrast to low-motility TAMΦs, TAMos feature a dramatically enhanced migration capability that help drive effective tumor infiltration of monocytes for subsequent differentiation and enrichment in TAMΦs in the tumor. Here, we draw a critical distinction between the well-studied chemotaxis of monocytes (i.e. their directionality or “steering wheel”) and their basal (random) migration (i.e. their net motion or “engine”), which functionally supersedes chemotaxis and are regulated by different molecular pathways. IL-6 secreted both by cancer cells and TAMos themselves is responsible for the high migration of TAMos by increasing their dendritic protrusion dynamics and myosin-based contractility via the JAK2/STAT3 signaling pathway. Moreover, TAMos - not TAMΦs - promote the proliferation of breast cancer cells through activation of the MAPK pathway. This work demonstrates the critical role that (random) migration plays in monocyte-driven TAMΦ enrichment in tumor. Based on these preclinical results, our work pinpoints IL-6 as a potential therapeutic target in combination with CCL2 to ameliorate current strategies against TAMΦ infiltration.

Considering the broad spectrum of pro-tumoral effects exerted by TAMΦs in the tumor microenvironment, inhibiting TAMΦ recruitment via chemotaxis as an anticancer therapy alone or in combination with chemotherapy and immunotherapy have been extensively explored in recent years. Nevertheless, it falls short in comparison with TAMΦ reprogramming strategies^67^ in that new studies and clinical trials have shifted towards anti-CD47/SIRPα antibody^68,69^, anti- CSF1/CSF1R antibody^70,71^, CD40 antagonist,^72^ PI3K inhibitor^73^, to name a few. Our work elucidates the limited efficacy of CCL2-based chemotaxis blockade in the syngeneic mouse study and identifies IL-6 as a potential therapeutic target that can exhibit a synergistic effect combined with CCL2 to further block monocyte-driven TAMΦ enrichment by targeting at monocyte random migration, which is independent of and supersedes its chemotaxis.

Although monocytes derived from bone marrow and recruited into the TME eventually differentiate into macrophages, the dramatic qualitative and quantitative differences in migration between monocytes and different phenotypes of macrophage, along with the unique pro-proliferation effect from TAMos instead of TAMΦs or naïve M0 macrophages established in this work imply that they are distinct populations of immune cells at the functional level. The near-complete loss of fast random migration capability in macrophages - featuring an 8-fold decreased speed and 16-fold decreased MSD - indicates that active infiltration of macrophages in TME is highly unlikely. Hence, we use the nomenclature “enrichment” instead of “infiltration” for macrophages. This enrichment is induced by the infiltration of highly migratory TAMos, which then differentiate into TAMΦs.

Standard *in vitro* culture of human primary macrophages, though optimized for differentiation, requires 5-7 days of M-CSF treatment for a complete differentiation of isolated monocytes into naïve M0 macrophages, followed by tumor conditioned medium re-education or cytokine combination to polarize them into terminal activated phenotypes^46,74^. In contrast, high-motility TAMos are induced within 6 h, indicating a fast functional phenotypic switch evidenced by IL-6R’s high expression on classical (CD14^+^CD16^-^) monocyte isolated directly from peripheral blood. Consequently, our work emphasizes the importance of pre-differentiated “tissue” monocytes at the early stage of recruitment, which is in alignment with the existence of various monocyte subsets with distinctive gene signatures compared with macrophages in tumor microenvironment via single-cell sequencing^51^. Unlike their well-characterized transmigration across the endothelium^75–80^, the migration of monocytes in tissue has attracted less attention, as they are typically viewed as macrophages that quickly differentiated once extravasated to replenish the tissue macrophage pool^18,81^. Limited studies on monocyte 3D migration either used monocytic cell lines^82^ or focused more on mechanisms driving monocyte’s ameboid-like migration^83–85^, but none incorporated the involvement of the tumor microenvironment in modulating their migration and how their migration compared to the migration of macrophages.

Although the surface marker expression^86–89^, phagocytotic capability^90^ and secretomics^91,92^ of macrophages activated with IL-4/13 share similarities with those of TAMΦs, our work shows that TAMΦs display a unique migratory phenotype of decreased motility compared to IL-4/13 activated macrophage and other macrophage phenotypes. In contrast, pro-inflammatory and regulatory macrophages induced by different combinations of cytokines differ significantly in marker expression and morphology, yet their migration resemble each other (Fig.1B-D). Correspondingly, emerging studies recently on single-cell analysis of macrophage populations in the tumor microenvironments^93–95^ have revealed unprecedented heterogeneity in mononuclear phagocyte clusters that is well beyond the simplistic bipolar M1/M2 classification system. This urges the addition of cell migration as a key functional characteristic in defining macrophage phenotype^96,97^. Future studies that seek to identify markers of high motility to facilitate the study of these subtypes in tissues will need to be undertaken.

The tumor microenvironment is a complex and continuously evolving entity in which tumor cells, stromal cells^98,99^ and immune cells interact with each other. In this study, IL-6 plays the central role of modulating TAMos’ high motility. Although we showed that IL-6 in our experimental systems was predominantly produced by cancer cells and TAMos themselves, it is possible that cancer-associated fibroblasts (CAFs)^100–102^, endothelial cells^103,104^ also contribute to the IL-6 reservoir. Nonetheless, IL-6 deficient TNBC tumors displayed significantly inhibited monocyte/macrophage infiltration. This to some extent demonstrates that tumor cells may function as the main source of IL-6 during TNBC tumor progression. Presently, only two monoclonal antibodies, namely tocilizumab (anti-IL-6R) and siltuximab (anti-IL-6) have been approved for treatment of rheumatoid arthritis and Castleman’s disease, but not for solid tumors^105^. Accumulated knowledge on IL-6’s critical role in breast tumor metastasis^106,107^, together with its induction of high-motility TAMos discovered in this work, urges clinical trials on blocking IL-6 in TNBC.

## Acknowledgments

The authors would like to thank all members of the Wirtz lab for their feedback. This work was partially supported through grants to D.W. from the National Cancer Institute (U54CA143868 and U54CA268083), the National Institute of Arthritis and Musculoskeletal and Skin Diseases (U54AR081774), and the National Institute on Aging (U01AG060903). All cartoons were created with BioRender.com.

## Author contributions

D.W. and W.D. conceptualized this project. W.D. designed and carried out most experiments. D.S. performed single cell analysis. A.C., B.Z. helped with mouse study. B.Z., A.T. helped with cell tracking. S.S. performed IMC. F.W. performed IF on spheroid sections. A.F., A.K. quantified IHC results with customized code. H. F. helped set up lattice light sheet microscopy. P.N., A.J. P.W., J.P provided customized MATLAB code for migration analysis. D.K. performed proteomics. The manuscript was written by W.D. with edits and input from D.W. and D.S.

## Methods

### PBMC isolation

Primary human monocytes were isolated and purified from peripheral blood mononuclear cells (PBMC) of leukopaks obtained from healthy volunteers (Blood donation center, Anne Arundel Medical Center, Annapolis, MD, USA) using Ficoll-Paque Plus (cytiva, 17144003) according to the manufacturer’s instructions. Briefly, 5 mL of blood was diluted in 30 mL RPMI 1640 (Gibco, 11875-093), and 13 mL of Ficoll-Paque Plus was slowly pipetted underneath the blood/RPMI mixture without disturbing the layer interface. Plasma, PBMCs, Ficoll-Paque Plus, and red blood cell layers were separated after centrifugation at 400 x g for 30 min with brake turned off. The PBMC layer was harvested, combined, and centrifuged (10 min, 400 x g). PBMCs were counted using 3% acetic acid with methylene blue (Stemcell, 07060) and were resuspended in freezing medium (90% FBS [Corning, 35-010-CV] + 10% DMSO [ATCC, 4-X]) at a concentration of 5 x 10^7^ cells/mL. Cryovials containing PBMCs were initially frozen at -80°C and transferred into liquid nitrogen for long-term storage.

### Monocyte isolation and macrophage differentiation

Macrophage culture medium was prepared using DMEM (Corning, 10-013-CV) supplemented with 10% heat-inactivated (57°C for 30 min) FBS (Corning, 35-010-CV), 1% penicillin-streptomycin (Sigma, P0781) and 50 ng/mL recombinant human M-CSF (R&D Systems, 216-MC). M1 macrophage culture medium was prepared using DMEM supplemented with 10% heat-inactivated FBS, 1% penicillin-streptomycin and 50 ng/mL recombinant human GM-CSF (R&D Systems, 215-MC). M1 macrophage differentiation medium was prepared using M1 macrophage culture medium supplemented with 100 ng/mL of LPS (Sigma-Aldrich, L2630) and 50 ng/mL of IFN-γ (R&D Systems, 285-IF). M2 macrophage differentiation medium was prepared using macrophage culture medium supplemented with 50 ng/mL IL4 (R&D Systems, 204-IL) and 50 ng/mL IL13 (R&D Systems, 213-ILB). Conditioned medium harvested from MDA-MB-231 organoids was used as tumor-associated macrophage (TAM) differentiation medium.

Primary human monocytes were isolated from PBMCs using negative selection human classical monocyte isolation kit (Miltenyi Biotec, 130-117-337) according to the manufacturer’s protocol. Isolated monocytes were counted using 3% acetic acid with methylene blue solution (Stemcell, 07060). Monocytes were seeded in a 6-well tissue culture plate (Corning, 353046) at 2.5 x 10^5^ cells/mL in macrophage culture medium. After 3 days, monocytes that remained in the suspension were gently aspirated away, the attached macrophages were washed three times with DPBS (Corning, 21-031-CV). For M0, M2 and Tumor-associated macrophages, fresh macrophage culture medium was added. Alternatively, for M1 macrophages, fresh M1 macrophage culture medium was added. On day 5, macrophages were washed with DPBS 3 times and corresponding differentiation media for different phenotypes was added. After 48 hours of incubation, macrophages were treated with pre-warmed TrypLE^TM^ Express Enzyme (Gibco, 12604013) for 10-15 min and were gently detached by pipetting.

### 3D collagen I gel preparation and cell random migration assay

3D collagen I gel embedded with cells were prepared as described previously by Hasini et al. (2017). Briefly, while keeping all components on ice, cell suspensions containing required number of cells in 1:1 (v/v) ratio of cell culture medium and reconstitution buffer (220 mg sodium bicarbonate [Sigma S5761] and 480 mg HEPES [Sigma H4034] dissolved in 10mL of ultrapure H2O, filtered with 0.22μm PES filter [Genesee 25-244]) were mixed with high concentration rat tail collagen I (Corning, 354249) to obtain a final concentration of 1 mg/mL. A calculated amount of 1M NaOH was quickly added, and the final mixture was thoroughly mixed to bring the pH to physiological level. 500 μL of cold, unsolidified gel mixture was added to each well of a 24-well tissue culture plate that was pre-warmed on a plate heater at 37°C. After 10-min of incubation on plate heater, the plate was transferred to incubator for an extra incubation of 1 hour to allow full polymerization before 500 μL medium containing cytokines/chemokines or conditioned medium was added on top of each gel. Unless specified otherwise, cells were seeded in gel at a standard density of 100 cells/mm^3^.

The 24-well plate was placed in live-cell box (Pathology Devices Inc.) that is maintained at 37°C/ 5% CO2 and was imaged via Nikon Eclipse TE-2000-E or Nikon Eclipse Ti2 system. Phase-contrast images were captured via a 10x objective at a time interval of 3 min for a total of 6 hours. Cells were tracked using MetaMorph (Molecular Devices)’s object tracking function. Multiple fields of views were set for each gel and a minimum of 100 individual cells were tracked per condition.

APRW model-based migration parameters including diffusivity, persistence time and anisotropy were computed from custom MATLAB code reported previously by Wu et al. For non-model-based migration characterization, time-independent cell speed was calculated as follows:

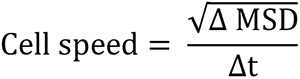

where ΔMSD is the difference in mean square displacement between 2 time points with an interval of Δt, which is half of the tracking time. For a total of N time points imaged, N/2 cell speeds (one each for the speed between time points ti and tN/2+i; i ranging from 1 to N/2) were calculated and were then averaged to attain the reported speed for one tracked single cell.

### Multi-compartment tumor-immune cell co-culture systems

Oil columns were prepared by adding 10 μL of serum-free DMEM medium and 50 μL mineral oil (Sigma, 69794) into a non-filter p10 tip (TipOne, 1111-3800). Matrigel matrix was diluted 1:1 v/v with DMEM medium on ice before usage. Following that, 50,000 MDA-MB-231 cancer cells were suspended in 1 μL of diluted Matrigel and pipetted into mineral oil to form a sphere that mimics solid MDA tumor core. Oil columns containing solid tumor cores were incubated at 37°C / 5% CO2 for 1 hour for further polymerization of Matrigel. MDA tumor cores were collected and washed in DMEM/DPBS (1:1 v/v mixture) to wash off remaining mineral oil.

For luciferase assay and monocyte tracking, one MDA tumor core was embedded in 100 μL of 1 mg/mL collagen I gel with desired density of monocytes/macrophages as described in abovementioned method section per well of a 96-well plate (Cellvis, P96-1.5H-N). 100 μL of culture medium was added on top of each gel after 1 hour incubation at 37°C / 5% CO2. For double-layer spheroid, one MDA tumor core was mixed with 10 μL of 1 mg/mL collagen I gel with desired density of monocytes/macrophages and pipetted into mineral oil to form a sphere. After 1 hour incubation at 37°C / 5% CO2, spheroids were harvested, washed and seeded into a round-bottom 96-well plate with 200 μL of DMEM. Phase-contrast images of spheroids were obtained using 4x objective.

### Flow cytometry

FACS wash buffer was made from DPBS (Gibco, 14190-144) containing 2% v/v FBS (Corning, 35-010-CV), 1mM EDTA (Invitrogen, 15575020), 0.1% w/v NaN3 (Sigma, S8032). Cell samples were washed 3 times in DPBS and resuspended at a concentration of 10^7^ cells/mL in 100 μL. Cell suspensions were blocked with 5 μL of Human TruStain FcX (Biolegend, 422301) for 15 min under room temperature. The fluorophore-conjugated antibodies (5 μL per antibody for 1 million cells) were then added into cell suspensions and incubated at 4 °C for 30 min, protected from light. Cells were then washed 3 times with 3 mL of FACS wash buffer and then resuspended in 500 μL of FACS wash buffer. Flow cytometry was carried out on a BD FACSCanto system and data analysis was performed using Flowjo Software version 10.4.

Antibodies used for labelling were as follows: APC anti-human CD14 (Biolegend, 367117), PE anti-human CD16 (Biolegend, 501106), PE/Cy7 anti-human CD126 (Biolegend, 352809), FITC anti-human CD181 (Biolegend, 320605), PE/Cyanine7 anti-human CD206 (Biolegend, 321124), APC anti-human CD80 (Biolegend, 375403), PE anti-human CD163 (Biolegend, 333605). Isotype control antibodies used for unstained samples were as follows: APC mouse IgG1 κ isotype ctrl (Biolegend, 400121), PE rat IgG1 κ isotype ctrl (Biolegend, 400407), PE/Cy7 mouse IgG1 κ isotype ctrl (Biolegend, 400125), FITC mouse IgG2b κ isotype ctrl (Biolegend, 400309).

### Western blot

3D collagen gels with monocytes were digested using 1:1 w/w collagenase type I (Gibco, 17100017) at 37°C / 5% CO2 for 45 min until no visible collagen fibers could be visualized. The lysate was filtered via a 40 μm nylon cell strainer (Falcon, 352340). The flow-through was centrifuged and washed in DPBS 3 times to yield a clean cell pellet.

Cell protein lysates were prepared via lysing cell pellets in clear sample buffer (0.5M Tris pH 6.8, 20% SDS, 50% glycerol in water) at 100°C for 5 min. Total protein concentration was evaluated using the Micro BCA^TM^ Protein Assay Kit (Thermo Scientific, 23235). Based on the calculation, 20 μg total proteins from different samples were loaded on 4-15% SDS-PAGE gel (Bio-Rad, 4561086) and ran in 1x Tris/Glycine/ SDS buffer (Bio-Rad, 1610772) at a voltage of 180V under room temperature for electrophoresis. Protein bands were transferred to PVDF membrane using the Trans-Blot Turbo system (Bio-Rad, 17001919). TBST buffer was made via supplementing 1x TBS buffer (Bio-Rad, 1706435) with 0.1% v/v TWEEN 20 (Sigma, P7949). The membrane was blocked in TBST with 5% BSA (Sigma, A9647) for 60 min at room temperature and incubated with primary antibodies at 4 °C for overnight. After 3 times wash in TBST, the membrane was incubated with secondary antibodies for 60 min at room temperature. After 3 times wash in TBST, 1 mL of ECL substrate solution (PROMETHEUS, 20-302) was added to the membrane and incubated for 2 min. Then the membrane was analyzed under the ChemiDoc^TM^ XRS+ imaging system (Bio-Rad). Images were analyzed using Image Lab software. Primary and secondary antibodies were used as follows: α-Tubulin (11H10) Rabbit mAb (CST, 2125), Arp2 (D85D5) Rabbit mAb (CST, 5614), phospho-myosin light chain 2 (Ser19) antibody (CST, 3671), Anti-Rabbit IgG HRP-linked Antibody (CST, 7074), IL6 (D5W4V) XP rabbit mAb (CST, 12912). Primary antibodies were diluted 1:1000 in TBST with 1% BSA and secondary antibody was diluted 1:5000 in TBST with 5% BSA for staining.

### Luciferase assay

Collagenase type I (Gibco, 17100017) solution was prepared at 2 mg/mL in DPBS. Culture medium from each gel was aspirated and each gel was washed 3 times with 100 μL of pre-warmed DPBS. 50 μL of collagenase solution was then added on top of each gel and incubated for 1 hour at 37°C / 5% CO2. Once the gel was digested into liquid form, 150 μL of BrightGlo- TritonX solution (90% BrightGlo [Promega, PRE2620] + 1% Triton X [Sigma, T9284]+ 9% DPBS) was applied on top of each gel. The plate was wrapped in aluminum foil and shaked at 400 rpm for 5 min, followed by 7-min sitting under room temperature. Bioluminescence was measured at a integration time of 500ms and 1000ms using a SpectraMax plate reader (Molecular devices). Specific proliferation was calculated using the following equation:

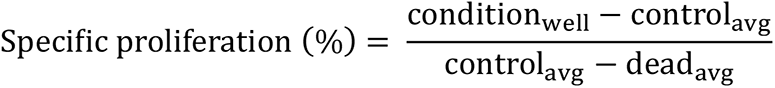

where dead condition wells were prepared via treating cells embedded in gel with 10% Triton X 1 hour prior to the start of collagenase digestion.

### Cell culture

MDA-MB-231 human triple-negative breast cancer cells (ATCC, HTB-26), 4T1 mouse breast cancer cells (ATCC, CRL-2539), THP-1 human monocytes (ATCC, TIB-202), U937 human monocytes (ATCC, CRL-1593.2), HEK-293T (ATCC, CRL-3216) were cultured according to ATCC recommendations in a humidified incubator maintained at 37°C and 5% CO2. MDA-MB-231 / HEK-293T cells were cultured in DMEM (Corning, 10-013-CV) supplemented with 10% FBS (Corning, 35-010-CV) and 1% penicillin-streptomycin (Sigma, P0781). 4T1 cells were cultured in RPMI 1640 (Gibco, 11875-093) supplemented with 10% FBS (Corning, 35-010-CV), 1% penicillin-streptomycin (Sigma, P0781), 1% HEPES (Gibco, 16530-080). THP-1, U937 monocytes were cultured in RPMI 1640 (Gibco, 11875-093) supplemented with 10% FBS (Corning, 35-010-CV), 1% penicillin-streptomycin (Sigma, P0781), 1% L-glutamine (Gibco, 25020-081), 1% HEPES (Gibco, 16530-080). All FBS added into medium was heat-inactivated at 37°C for 30 min. Luciferase-tagged MDA-MB-231 cells, Lifeact-GFP-tagged U937 cells were cultured in the same complete media as wild-type cells. IL6-KD 4T1 cells were cultured in wild-type complete medium supplemented with 5 μg/mL puromycin (SCBT, sc-108071).

### Lentivirus production and transduction

HEK-293T cells were seeded 24 hours before transfection at a density of 2.5x10^5^ in 2mL per well of a 6-well plate. Expression vector pLKV-EF1a-mCherryLuciferase-W (Addgene, 200101) or pLenti-Lifeact EGFP (Addgene, 187686) was mixed with psPAX2 (Addgene, 12260) and pMD2.G (Addgene, 12259) at a ratio of 3 μg:3 μg:1.5 μg and added to 16 μL of 2M CaCl2 solution. PCR water (Quality Biological, 351-161-671) was added directly to mixture to bring the final volume to 125 μL. After 5 min, 125 μL 2x HBS buffer was added and mixed well. Let the DNA/CaCl2 mixture sit at R.T. for 30 min and then add the mixture dropwise on top of 293T cells. Virus supernatant was harvested 48h post-transfection, filtered via 0.45μm PES filter (Genesee, 25-246) and stored under -80°C for future lentiviral transduction. For IL6 knock down of 4T1 cells, lentiviral particles were directly purchased: IL6 shRNA (m) lentiviral particles (SCBT, sc-39628-V), control shRNA lentiviral particles (SCBT, sc-108080). Lentivirus titer was determined via Lenti-X p24 rapid titer kit (Takara, 631476) per manufacturer’s protocol.

For transduction, cells were seeded at a density of 8x10^4^ per well of a 12-well plate and cultured till reaching a confluency of around 50%. Lentivirus particles were resuspended in cell complete culture medium supplemented with 5 μg/mL polybrene (SCBT, sc-134220) at a MOI of 1-10 and then added dropwise. The plate was swirled gently for mixing and incubated overnight at 37°C / 5% CO2. After that, medium with lentiviral particles and polybrene was aspirated and fresh complete medium was added. Cells were split when reaching 95% confluency and kept culturing. Transduced mCherry-positive MDA-MB-231 and EGFP-positive U937 cells were sorted by fluorescence-activated cell sorting (FACS, >95%) with a SH800S Cell Sorter (Sony Biotechnology). For IL6 knock down, 4T1 cells were subsequently cultured in complete medium with 5 μg/mL puromycin for selection (SCBT, sc-108071).

### Transwell assay

Low concentration collagen I (Corning, 354236) was diluted in 0.1% acetic acid (17.5 mM in DPBS) to a final concentration of 100 μg/mL. 32 μL of diluted collagen I was added to each Boyden chamber insert (Sigma, CLS3472) and the insert was left under R.T. for 1 hour for polymerization of collagen into a thin coating layer. Excessive liquid was gently aspirated off. After 3 times wash in DPBS, DMEM complete medium was added to the inset and bottom chamber for 1h pre- conditioning under 37°C / 5% CO2. 50,000 freshly isolated monocytes were resuspended in 200 μL of DMEM complete medium in the insert and corresponding conditioned media and cytokines / chemokines were added to the bottom chamber for initialization of cell transmigration assay. 16 hours post assay, monocytes transmigrated to the bottom chamber were counted via applying particle analysis function in ImageJ on 10x phase-contrast images taken at multiple fields of views in each well.

### Lattice light sheet microscopy

Lifeact-tagged U937 monocytic cells were embedded in 1 mg/mL collagen gel at a density of 100 cells/mm^3^ and mounted on a 5mm glass coverslip. Samples were sent and imaged using a 3i lattice light-sheet microscope by Johns Hopkins Medicine Microscope Facility.

### PrestoBlue cell viability assay

Organoids were incubated at 37 °C in a 1× PrestoBlue reagent (Invitrogen, A13261) diluted in complete DMEM for 3 h protected from light. RFUs were read with 540nm excitation at 600nm using SpectraMax plate reader (Molecular devices). After reading, organoids were wash in DBPS 3× and given 200 µL of fresh DMEM medium. The experiment was carried out for up to 4 days. To account for background signal, signals from 10 wells of 1× PrestoBlue were measured, averaged, and subtracted. The average background signal value was subtracted from the experimental wells.

### RNA extraction from 3D spheroids and RT-qPCR analysis

Total RNA from MDA-MB-231 cells embedded in spheroids was isolated using RNeasy Micro Kit (QIAGEN, 74004) per manufacture protocol. Extensive vortexing and mixing was performed for higher yield of RNA from 3D matrix. cDNA synthesis was performed using iScript cDNA Synthesis Kit (Bio-Rad, 1708890). Real-time PCR reactions were set up using iTaq Universal SYBR Green Supermix (Bio-Rad, 1725121) and were executed in a thermal cycler (CFX384^TM^ Real-Time System, Bio-Rad). The primers designed for specific gene amplification are listed in Table 1. Relative quantitation was performed using the △△Ct method in CFX Manager software.

**Table 1.**
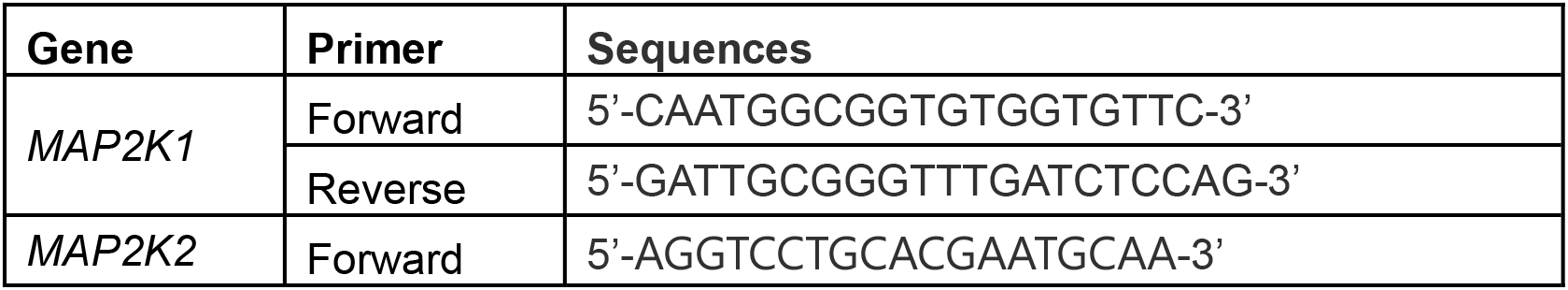
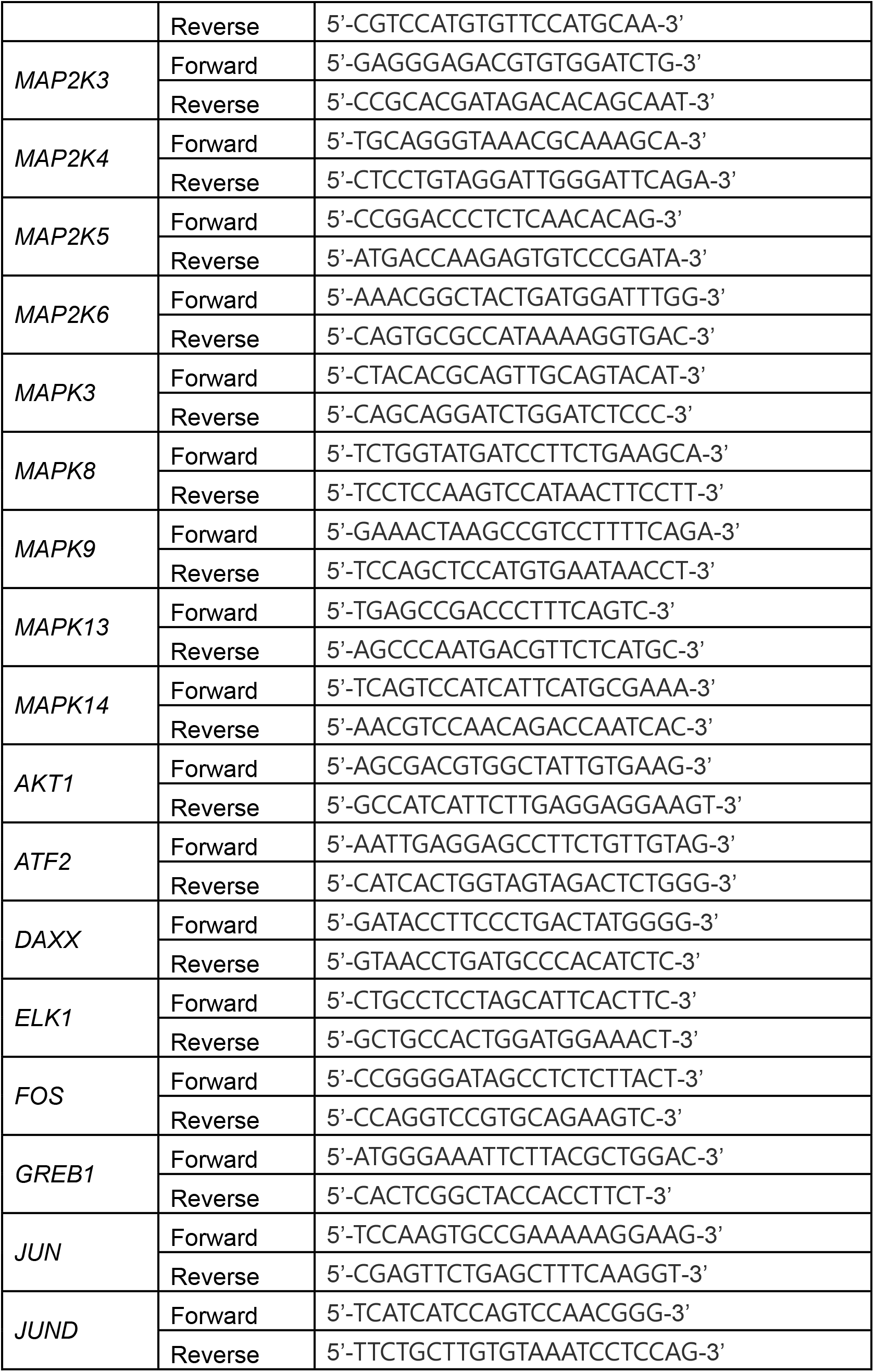
Primer pairs of genes associated with MAPK pathway

**Table 2.** Antibody list for imaging mass cytometry

| Mass | Metal | Antigen | Clone | Dilution Factor | Source |
| --- | --- | --- | --- | --- | --- |
| 141 | Pr | S100A9 | D3U8M | 250 | Cell Signaling Technology® |
| 142 | Nd | Ly-6C | EPR27220-23 | 250 | Abcam |
| 150 | Nd | Ki-67 | B56 | 250 | Standard BioTools™ |
| 156 | Gd | F4/80 | D2S9R | 250 | Standard BioTools™ |
| 159 | Tb | CD4 | BLR167J | 250 | Standard BioTools™ |
| 162 | Dy | CD8 | EPR21769 | 250 | Standard BioTools™ |
| 163 | Dy | CD11b | EPR1344 | 250 | Standard BioTools™ |
| 191 | Ir | DNA 1 |  |  | Standard BioTools™ |
| 193 | Ir | DNA 2 |  |  | Standard BioTools™ |
| 96-104 | Ru | Counterstain |  |  | Electron Microscopy Sciences |

### 4T1 syngeneic mouse study

All animal experiments were approved by the Institutional Animal Care and Use Committee (IACUC) of the Johns Hopkins University (protocol MO19E327). Mice were kept at 5 per cage for all groups. The study was performed using 6-8-week-old BALB/cJ (Jackson Laboratory, strain 000651) female mice. For all mice, fur around left mammary fat pad area was removed via Nair treatment one day before tumor cell injection. 2x10^4^ 4T1 cells were resuspended in 50 μL of ice-cold 1:1 DPBS / Matrigel solution and orthotopically injected into mammary fat pad using a 26- gauge syringe. After tumors became palpable at around day 8, mCCR2 antagonist INCB3344 was administered at 2 mg/kg twice a week via intraperitoneal injection. Tumors were measured using digital calipers approximately twice per week at indicated days and mice were weighed at the end of each experiment. Tumor volumes were calculated using the following formula: 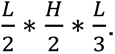 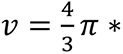 Tumors and organs were harvested at indicated days and were fixed in formalin and embedded in paraffin. FFPE sections were H&E stained and IHC stained for anti-mouse CD11b and CD45 to measure pan-monocyte/macrophage infiltration and common leukocytes infiltration by The Johns Hopkins Oncology Tissue Service core facility. All histological slides were scanned using NanoZoomer S210 (Hamamatsu) at 20x magnification.

### Imaging mass cytometry (IMC)

Slides were cut from the FFPE blocks and immunohistochemistry with mass cytometry antibodies was performed. The slides were baked for two hours, dewaxed in xylene, rehydrated in a descending alcohol gradient (100%, 95%, 80%, 70% EtOH in Maxpar® H2O), then washed with Maxpar® H2O. The slides then incubated in Cell Conditioning Solution (Roche PN 5279801001) at a sub-boiling temperature for 1 hour. Slides were blocked with 3% BSA in Maxpar PBS for 45 minutes at room temperature. An imaging mass cytometry antibody cocktail – listed in Table# - stained the slides overnight at 4 degrees. Custom antibodies were conjugated in-house, diluted to a concentration of 0.25 mg/mL to 0.5 mg/mL, then titrated empirically. For DNA labelling, Cell- ID™ Intercalator-Ir (Standard Biotools PN 201192A) was diluted at 1:400 in Maxpar® PBS. As a counterstain, Ruthenium tetroxide 0.5% Aqueous Solution (Electron Microscopy Sciences PN 20700-05) was diluted at 1:2000 in Maxpar® PBS. Images were acquired with a Hyperion Imaging System (Standard BioTools at the Johns Hopkins Mass Cytometry Facility. Upon acquisition, representative images were through MCD Viewer™ (Standard BioTools).

### Immunohistochemistry quantifications

Histological slides were scanned using a Hamamatsu S210 scanner at 20x resolution (0.5 µm/pixel). Scanned images subsequently downsampled to 10x magnification (1 µm/pixel) and saved as tiff image files. Color deconvolution was applied to isolate the CD45-positive immunohistochemical signal into a separate image channel. Within each isolated image channel, intensity peaks were identified, and spatial coordinates were collected (x, and y). Assessment of tumor CD45+ cell density was performed by normalizing the cell count by total tissue area of each individual tumor.

### Statistical analysis

Graphpad Prism 10 software was used for statistical analysis. An unpaired student’s t-test with Welch’s correction or nonparametric Mann-Whitney U test was performed to evaluate the statistical significance between two groups. For column analysis of multiple conditions, Kruskal-Wallis with Dunn’s multiple comparison test was performed. Significant values were given in grades ns P˃0.05, *P≤0.05, **P≤0.01, ***P≤0.001, ****P≤0.0001.

