## Supplementary Figures for "High-motility pro-tumorigenic monocytes drive macrophage enrichment in the tumor microenvironment"

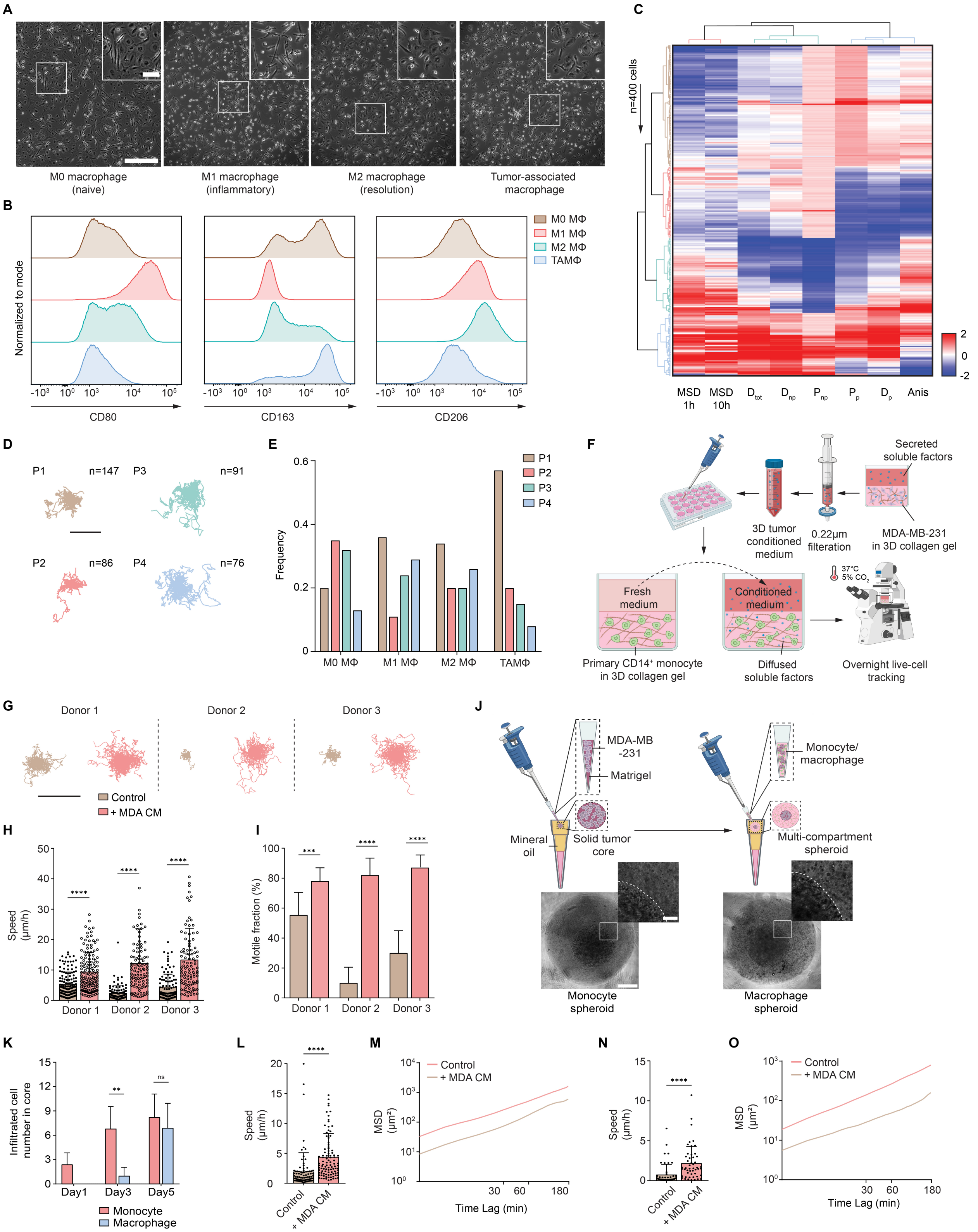
A
M0 macrophage (naive)
M1 macrophage (inflammatory)
M2 macrophage (resolution)
Tumor-associated macrophage
B
Normalized to mode
CD80
CD163
CD206
M0 MΦ
M1 MΦ
M2 MΦ
TAMΦ
C
n=400 cells
MSD 1h
MSD 10h
D tot
D np
P np
P p
D p
Anis
2
0
-2
D
P1 n=147
P3 n=91
P2 n=86
P4 n=76
E
Frequency
P1
P2
P3
P4
M0 MΦ
M1 MΦ
M2 MΦ
TAMΦ
F
Secreted soluble factors
3D tumor conditioned medium
0.22μm filteration
MDA-MB-231 in 3D collagen gel
Fresh medium
Conditioned medium
37°C 5% CO2
Primary CD14+ monocyte in 3D collagen gel
Diffused soluble factors
Overnight live-cell tracking
G
Donor 1
Donor 2
Donor 3
Control
+ MDA CM
H
Speed (μm/h)
Donor 1
Donor 2
Donor 3
I
Motile fraction (%)
Donor 1
Donor 2
Donor 3
J
MDA-MB -231
Matrigel
Mineral oil
Solid tumor core
Monocyte/ macrophage
Multi-compartment spheroid
Monocyte spheroid
Macrophage spheroid
K
Infiltrated cell number in core
Day1
Day3
Day5
Monocyte
Macrophage
L
Speed (μm/h)
Control
+ MDA CM
M
MSD (μm²)
Control
+ MDA CM
Time Lag (min)
N
Speed (μm/h)
Control
+ MDA CM
O
MSD (μm²)
Control
+ MDA CM
Time Lag (min)

**Supplementary Fig. S1** (**A**) Representative phase contrast images taken with a 20x objective for different phenotypes of macrophages. Scale bar, 200 μm (inset, scale bar, 50 μm). (**B**) CD80 (M1 marker), CD163 (M2/TAMΦ marker), CD206 (M2 marker) expressions on different phenotypes of macrophages. Counts were normalized to mode for each group. (**C**) Heatmap showing the magnitude of 8 macrophage migration parameters per cell (n = 400 cells, 100 cells per phenotype of macrophage) including MSD at 1 h, MSD at 10h, diffusivity on primary axis ($D_p$), persistence on primary axis ($P_p$), diffusivity on secondary axis ($D_{np}$), persistence on secondary axis ($P_{np}$), total diffusivity ($D_{tot}$), anisotropy (Phi). Each row represents a single cell, and each column represents one parameter. 4 spatio-temporal clusters were classified (P1-P4). Cityblock was used as the distance metric and ward was used as linkage metric. (**D**) Trajectories of all cells in each cluster. Scale bar, 150 μm. (**E**) Frequencies of each spatio-temporal cluster in different phenotypes of macrophages. (**F**) Illustration cartoon for monocyte migration assay. 3D Conditioned medium from MDA-MB-231 cells embedded in collagen matrix were harvested and applied on monocytes. Tracking was initiated after 6 h of incubation under 37 °C, 5% $CO_2$. (**G-I**) Trajectories, speeds and motile fractions of monocytes isolated from 3 independent donors upon treatment of MDA-MB-231 conditioned medium. Motile monocytes were defined as migrating more than their own size within 1 h ($MSD>D^2$ = 100 $\mu m^2$, where D = 10 μm is the average monocyte diameter). Scale bar, 250 μm. n = 135 individual cells tracked for donor 1. n = 100 individual cells tracked for donors 2 and 3. (**J**) Illustration cartoon for multi-compartment spheroid. Representative phase contrast images taken with a 4x objective showed even distribution of monocytes and macrophages in the outer collagen layer on day 0. Scale bar, 300 μm (inset scale bar, 30 μm). (**K**) Quantification of infiltrated cell numbers in monocyte and macrophage spheroids on day1, day 3 and day 5. n = 10 technical replicates. Data are plotted as mean + standard deviation. Unpaired t test with Welch's correction was used for statistical analysis (**P≤0.01). (**L**) Characterized speeds of THP-1 cells with or without MDA CM treatment. (**M**) Mean MSD of all tracked THP-1 cells plotted against time lag (min). (**N**) Characterized speeds of U937 cells with or without MDA CM treatment. (**O**) Mean MSD of all tracked U937 cells plotted against time lag (min). For migration assays, characterized speeds are plotted as mean + standard deviation. Non-parametric Mann-Whitney test was used for statistical analysis (****P≤0.0001, ***P≤0.001).

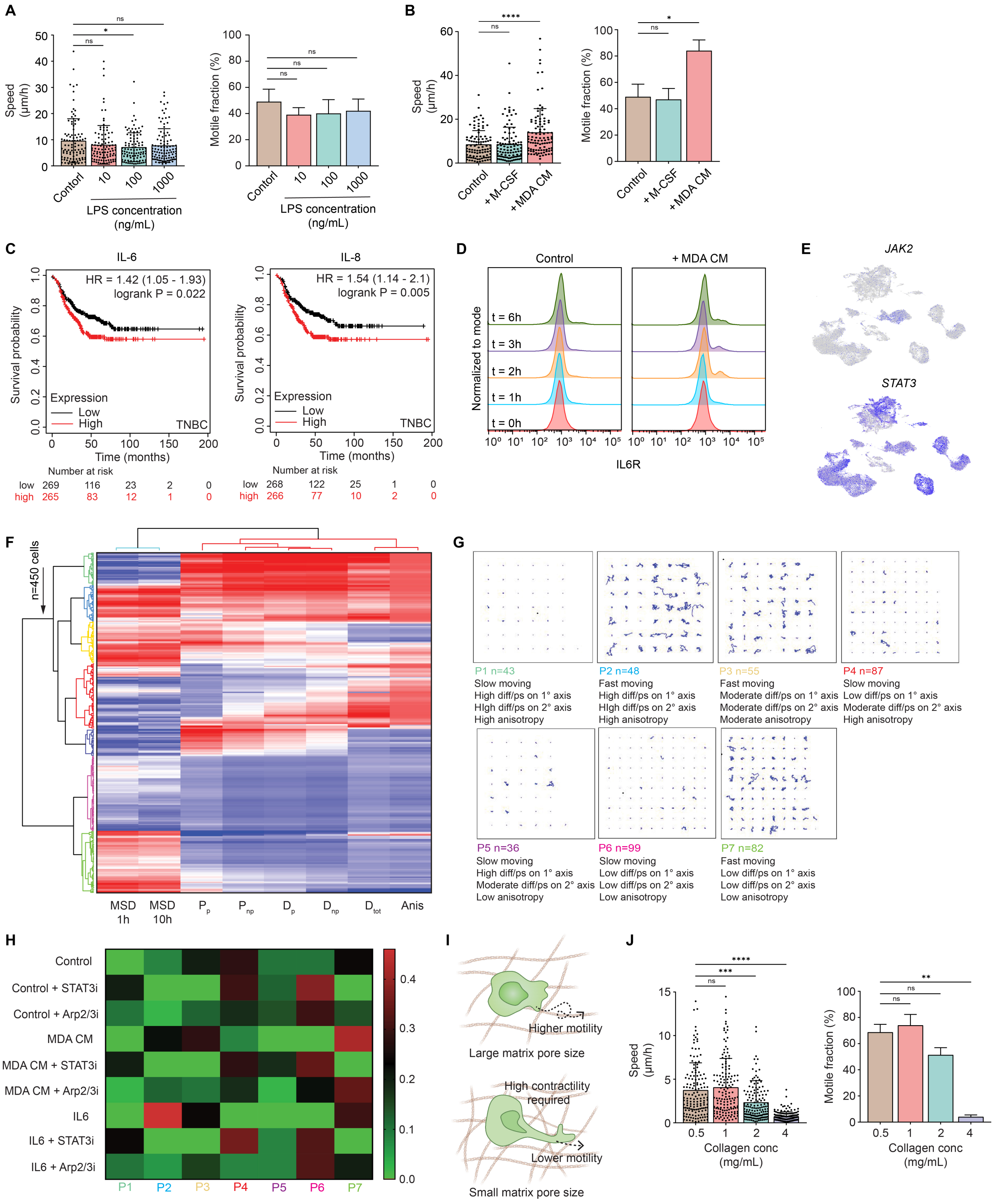
A
Speed (μm/h)
ns
*
ns
Contorl
10
100
1000
LPS concentration (ng/mL)
Motile fraction (%)
ns
ns
ns
B
Speed (μm/h)
****
ns
Control
+ M-CSF
+ MDA CM
Motile fraction (%)
*
ns
C
IL-6
HR = 1.42 (1.05 - 1.93)
logrank P = 0.022
Survival probability
Expression
Low
High
TNBC
Time (months)
Number at risk
low 269 116 23 2 0
high 265 83 12 1 0
IL-8
HR = 1.54 (1.14 - 2.1)
logrank P = 0.005
Survival probability
Expression
Low
High
TNBC
Time (months)
Number at risk
low 268 122 25 1 0
high 266 77 10 2 0
D
Control
+ MDA CM
Normalized to mode
t = 6h
t = 3h
t = 2h
t = 1h
t = 0h
IL6R
E
JAK2
STAT3
F
n=450 cells
MSD 1h
MSD 10h
Pp
Pnp
Dp
Dnp
Dtot
Anis
G
P1 n=43
Slow moving
High diff/ps on 1° axis
High diff/ps on 2° axis
High anisotropy
P2 n=48
Fast moving
High diff/ps on 1° axis
High diff/ps on 2° axis
High anisotropy
P3 n=55
Fast moving
Moderate diff/ps on 1° axis
Moderate diff/ps on 2° axis
Moderate anisotropy
P4 n=87
Slow moving
Low diff/ps on 1° axis
Moderate diff/ps on 2° axis
High anisotropy
P5 n=36
Slow moving
High diff/ps on 1° axis
Moderate diff/ps on 2° axis
Low anisotropy
P6 n=99
Slow moving
Low diff/ps on 1° axis
Low diff/ps on 2° axis
Low anisotropy
P7 n=82
Fast moving
Low diff/ps on 1° axis
Low diff/ps on 2° axis
Low anisotropy
H
Control
Control + STAT3i
Control + Arp2/3i
MDA CM
MDA CM + STAT3i
MDA CM + Arp2/3i
IL6
IL6 + STAT3i
IL6 + Arp2/3i
P1
P2
P3
P4
P5
P6
P7
I
Higher motility
Large matrix pore size
High contractility required
Lower motility
Small matrix pore size
J
Speed (μm/h)
****
***
ns
Collagen conc (mg/mL)
Motile fraction (%)
**
ns
ns
Collagen conc (mg/mL)

**Supplementary Fig. S2** (**A**) Characterized speeds and motile fractions of monocytes activated with a titration of LPS (10/100/1000 ng/mL). (**B**) Characterized speeds and motile fractions of monocytes treated with 50 ng/mL M-CSF and MDA CM. (**C**) Kaplan-Meier recurrence-free survival plots for TNBC subtype (534 patients evaluated), grouped based on IL-6 or IL-8 expression. A median split was used to define high versus low IL-6 or IL-8 expression. HR, hazard ratio. (**D**) Short-term IL-6R expression on monocytes treated with MDA CM. Counts were normalized to mode for different time points. (**E**) *JAK2* and *STAT3* expression in single cell atlas of human breast cancer (high – blue; low – gray). (**F**) Heatmap showing the magnitude of 8 migration parameters per monocyte (n = 450 cells, 50 cells per condition). Parameters presented are the same as described in Fig. S1C. 7 spatio-temporal clusters were classified (P1-P7). Cityblock was used as the distance metric and ward was used as linkage metric. (**G**) Trajectories of all cells and migration features of each cluster. (**H**) Heatmap of each spatio-temporal cluster's frequency in different conditions. (**I**) Illustration cartoon of monocyte migrating in 3D collagen matrices with large or small pore sizes. (**J**) Characterized speeds and motile fractions of monocytes embedded in 3D matrices with increased collagen concentration. Time-independent speeds (N = 2 donors, n = 50 cells for a total of 100 individual cells tracked) and motile fractions (N = 2 donors, n = 5 technical replicates) are plotted as mean + standard deviation. Kruskal-Wallis ANOVA with Dunn's test was used for statistical analysis comparing all groups with control (****$P \leq 0.0001$, ***$P \leq 0.001$, *$P \leq 0.05$, ns $P > 0.05$).

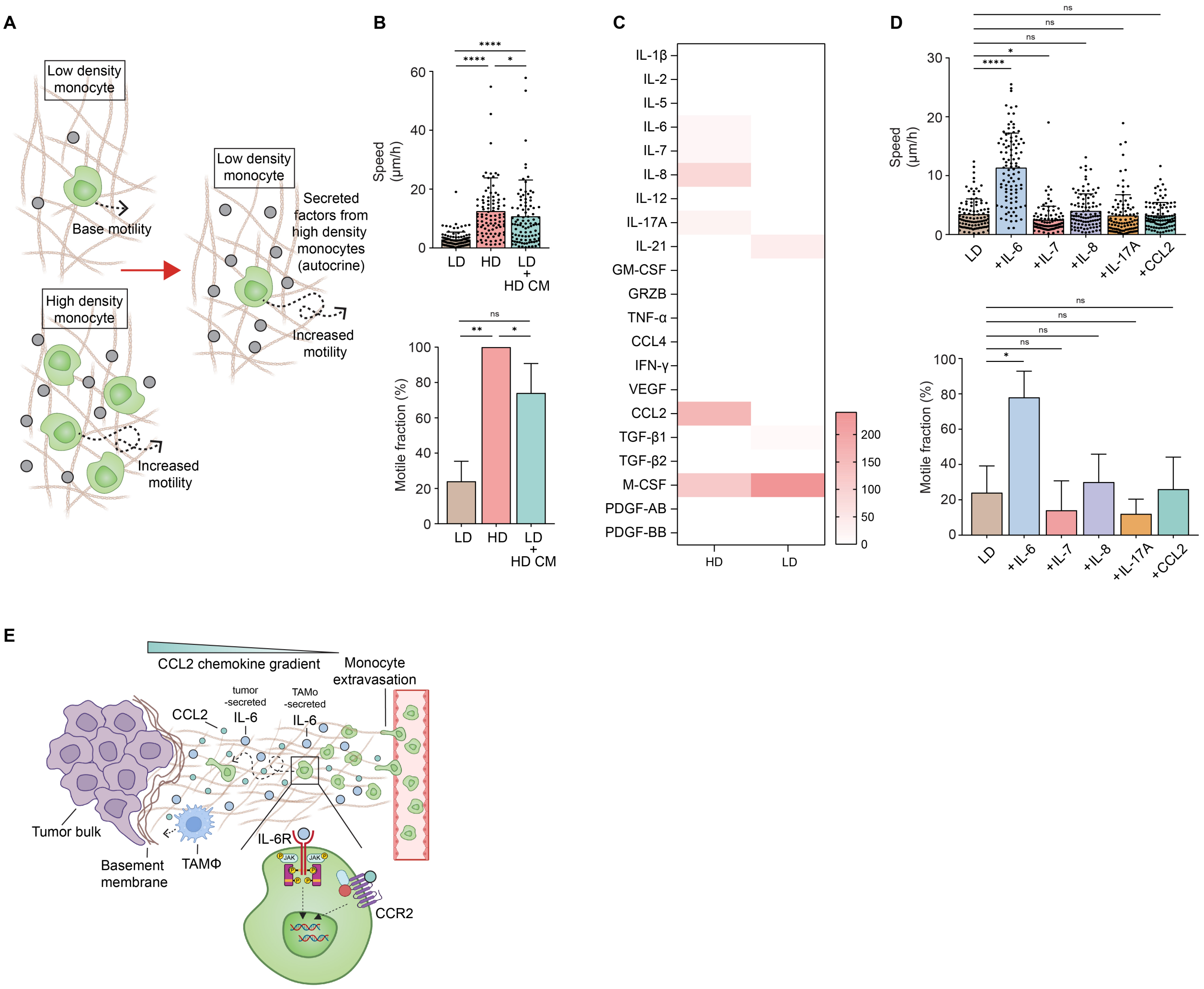
A
Low density monocyte
Base motility
High density monocyte
Increased motility
Low density monocyte
Secreted factors from high density monocytes (autocrine)
Increased motility
B
Speed (μm/h)
LD
HD
LD + HD CM
Motile fraction (%)
C
IL-1β
IL-2
IL-5
IL-6
IL-7
IL-8
IL-12
IL-17A
IL-21
GM-CSF
GRZB
TNF-α
CCL4
IFN-γ
VEGF
CCL2
TGF-β1
TGF-β2
M-CSF
PDGF-AB
PDGF-BB
HD
LD
D
Speed (μm/h)
LD
+IL-6
+IL-7
+IL-8
+IL-17A
+CCL2
Motile fraction (%)
E
CCL2 chemokine gradient
Monocyte extravasation
CCL2
tumor -secreted IL-6
TAMΦ -secreted IL-6
Tumor bulk
Basement membrane
TAMΦ
IL-6R
JAK
CCR2

**Supplementary Fig. S3** (**A**) HD monocytes (high density, 500 cells/mm$^3$) were incubated in 3D collagen matrix for 24 h. Supernatant from HD monocytes containing secreted factors were harvested and applied back on freshly seeded LD monocytes (100 cells/mm$^3$). (**B**) Characterized speeds and motile fractions of monocytes embedded at LD, HD and LD treated with HD conditioned medium. (**C**) Molecules secreted by LD and HD monocytes incubated in 3D collagen matrix for 48 h. Proteomics were run on supernatants to generate average intensity of a human immune cytokine panel using Isoplexis Isocode chips. (**D**) Characterized speeds and motile fractions of LD monocytes treated with molecules identified from proteomics assay. (**E**) Illustration cartoon demonstrating the central role of IL-6 in monocyte random migration. Monocytes recruited by CCL2 extravasated, induced by high concentration of IL-6 in the TME secreted by tumor core into high-motility functional phenotype and in the meanwhile secrete IL-6 to fuel themselves. IL-6 mediated enhanced random migration and CCL2 mediated chemotaxis then combine to drive the infiltration of monocyte. Recruited monocytes in tumor invasive margin and core are eventually differentiated into tumor-associated macrophages with low motility. For migration assays, characterized speeds (N = 2 donors, n = 50 cells for a total of 100 individual cells tracked) and motile fractions (N = 2 donors, n = 5 technical replicates) are plotted as mean + standard deviation. Non-parametric Mann-Whitney test was used for statistical analysis comparing 2 groups (****$P \leq 0.0001$, **$P \leq 0.01$, *$P \leq 0.05$, ns $P > 0.05$). Kruskal-Wallis ANOVA with Dunn's test was used for statistical analysis for multiple comparisons (****$P \leq 0.0001$, *$P \leq 0.05$, ns $P > 0.05$).

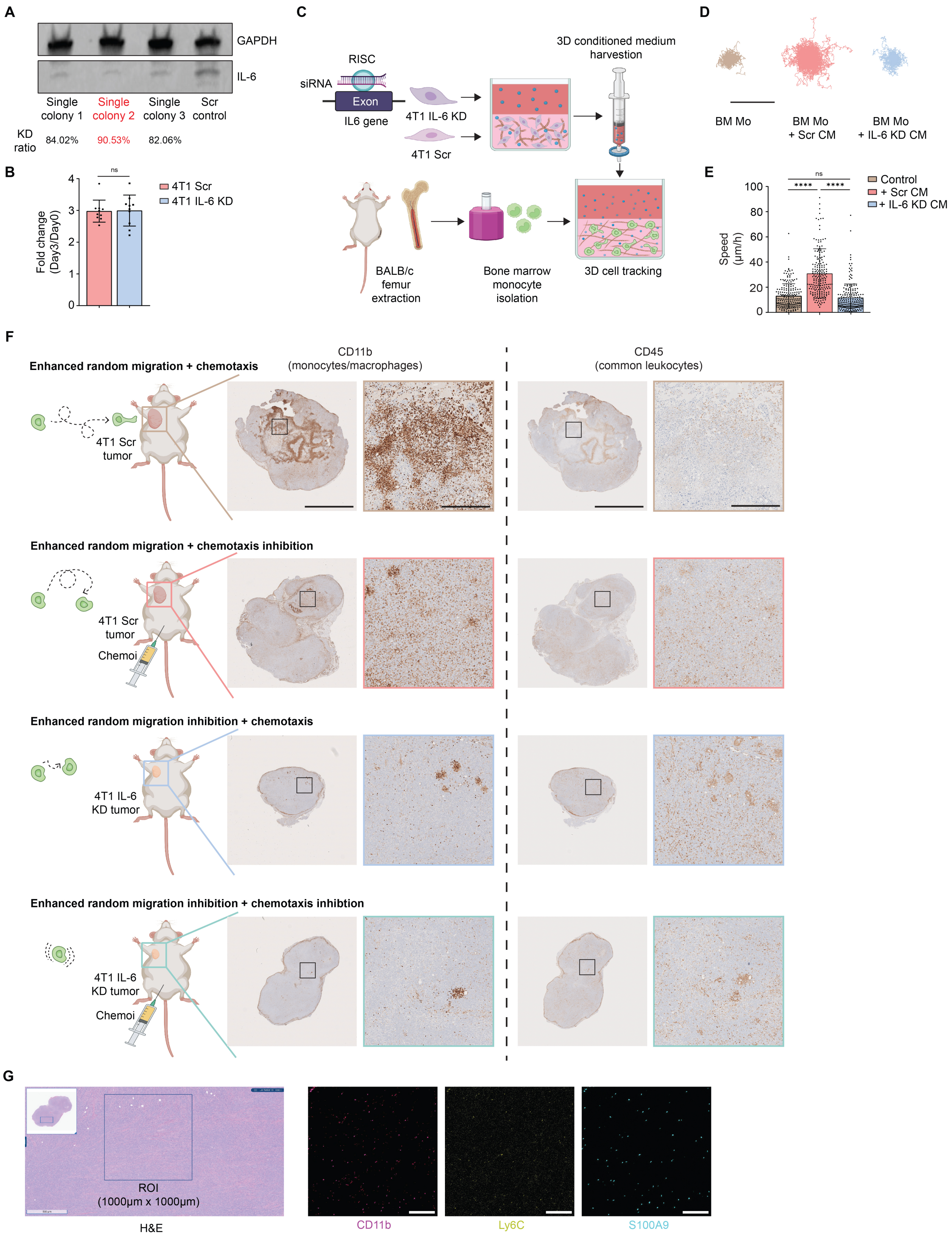
A
GAPDH
IL-6
Single colony 1
Single colony 2
Single colony 3
Scr control
KD ratio
84.02%
90.53%
82.06%
B
Fold change (Day3/Day0)
ns
4T1 Scr
4T1 IL-6 KD
C
RISC
siRNA
Exon
IL6 gene
4T1 IL-6 KD
4T1 Scr
3D conditioned medium harvestion
BALB/c femur extraction
Bone marrow monocyte isolation
3D cell tracking
D
BM Mo
BM Mo + Scr CM
BM Mo + IL-6 KD CM
E
Speed (μm/h)
ns
****
****
Control
+ Scr CM
+ IL-6 KD CM
F
CD11b (monocytes/macrophages)
CD45 (common leukocytes)
Enhanced random migration + chemotaxis
4T1 Scr tumor
Enhanced random migration + chemotaxis inhibition
4T1 Scr tumor
Chemoi
Enhanced random migration inhibition + chemotaxis
4T1 IL-6 KD tumor
Enhanced random migration inhibition + chemotaxis inhibtion
4T1 IL-6 KD tumor
Chemoi
G
ROI (1000μm x 1000μm)
H&E
CD11b
Ly6C
S100A9

**Supplementary Fig. S4** (**A**) Western blot validating the successfully knock down of IL-6 in 4T1 cells. (**B**) PrestoBlue signal fold change of 4T1 spheroids after 72 h incubation. n=10 technical replicates. Unpaired t test with Welch's correction was used for statistical analysis (ns $P>0.05$). (**C**) Illustration cartoon of mouse bone marrow (BM) monocyte migration assay. 3D conditioned media from 4T1 Scr and IL-6 KD cells were harvested and applied on mouse BM monocytes (isolated from mouse femur bone marrow via negative selection) embedded in 3D collagen matrix. (**D**) Plotted trajectories of mouse BM monocytes. Scale bar, 500 µm. (**E**) Characterized speeds of mouse BM monocytes. Non-parametric Mann-Whitney test was used for statistical analysis comparing Scr CM and IL-6 KD CM conditions. (****$P\leq0.0001$). Kruskal-Wallis ANOVA with Dunn's test was used for statistical analysis comparing all conditions with control (****$P\leq0.0001$, ns $P>0.05$). (**F**) Representative scanned IHC images stained with CD11b and CD45 of mouse tumors from different groups. Scale bar, 5mm (ROI scale bar, 500 µm). (**G**) Representative H&E and IMC images of one ROI in the group with enhanced migration and chemotaxis inhibition. For IMC images: CD11b-magenta channel, Ly6C-yellow channel, S100A9-cyan channel. Scale bar, 200 µm.

**Supplementary Video 1**: Tracking videos of monocytes and macrophages upon long term MDA CM treatment. Monocyte migration was monitored on day 1 and macrophage migration was monitored on day 7 with complete differentiation. Cells were tracked for 12 h with a time interval of 3 min. Scale bar, 200 μm.

**Supplementary Video 2**: Representative sub-minute tracking of monocyte's dendritic protrusion dynamics via lattice light sheet microscopy. Videos were taken for 11 min and 10 sec with a time interval of 10 sec. Scale bar, 5 μm.
